# Desert Hedgehog mediates stem Leydig cell differentiation through Ptch2/Gli1/Sf1 signaling axis

**DOI:** 10.1101/2025.06.13.659479

**Authors:** Changle Zhao, Yongxun Chen, Lei Liu, Xiang Liu, Hesheng Xiao, Feilong Wang, Qin Huang, Xiangyan Dai, Wenjing Tao, Deshou Wang, Jing Wei

## Abstract

Desert Hedgehog (Dhh) mutations cause Leydig cell dysfunction, yet the mechanisms governing Leydig lineage commitment through Dhh-mediated receptor selectivity, transcriptional effector specificity, and steroidogenic coupling remain elusive. In this study, using CRISPR/Cas9-mediated gene knockout and stem Leydig cells (SLCs) transplantation, we identified a critical Dhh/Patched 2 (Ptch2)/Glioma-associated oncogene homolog 1 (Gli1)/steroidogenic factor 1 (Sf1) signaling axis essential for SLC differentiation in Nile tilapia *(Oreochromis niloticus)*. Dhh deficiency resulted in defective adult Leydig cells and androgen insufficiency. Rescue experiments involving 11-ketotestosterone administration and a Dhh agonist treatment, combined with SLCs transplantation, demonstrated that Dhh regulates SLC differentiation, not survival. *In vitro* knockout of *ptch1* and *ptch2* in SLCs revealed that Ptch2 likely acts as the functional receptor for Dhh. This was further supported by *in vivo* genetic rescue experiments, where *ptch2* mutation did not impair testicular development, yet completely rescued the testicular defects in *dhh* mutants—consistent with Ptch2 acting as an inhibitory receptor whose loss alleviates Dhh pathway suppression. Luciferase assays in Gli-knockout SLCs demonstrated that Gli1 acts as the primary transcriptional effector, and transactivates *sf1* expression. Additionally, functional transplantation assays confirmed that Sf1 is indispensable for SLCs differentiation, as Sf1-overexpressing SLCs rescued differentiation, whereas *sf1*-mutant SLCs failed. Overall, our work delineates the Dhh-Ptch2-Gli1-Sf1 axis and provides fundamental insights into the endocrine regulation of Leydig cell lineage development.

**Highlights:**

- Dhh regulates the differentiation of SLCs rather than their survival
- Ptch2 acts as the functional receptor for Dhh signaling in SLCs
- Gli1 is the principal transcriptional activator for Dhh signaling in SLCs
- Sf1 is the critical downstream effector of Gli1 in SLCs differentiation

## INTRODUCTION

Leydig cells are the primary source of androgens, producing approximately 95% of circulating testosterone to sustain spermatogenesis and male fertility (Inoue, Baba, and Morohashi, 2018; Jiang et al., 2014). Leydig cells originate from stem Leydig cells (SLCs), which differentiate into steroidogenic adult Leydig cells (ALCs) under paracrine regulation (P. Chen, Zirkin, and Chen, 2020; S. Yao et al., 2022). Steroidogenic factor 1 (Sf1), an orphan nuclear receptor transcription factor, serves as a master regulator of this process by activating steroidogenic enzymes (e.g., cytochrome P450, family 11, subfamily C, polypeptide 1 [Cyp11c1], 3β-hydroxysteroid dehydrogenase [3β-HSD]) to drive ALC maturation (Parker et al., 2002). In mice, mutations of *Sf1* exhibit arrested testicular development and abolished steroidogenesis (Houzelstein et al., 2024; Park, Tong, and Jameson, 2007; Smith et al., 2023), while human *SF1* mutations are associated with 46,XY disorders of sexual development (DSD), such as Swyer syndrome (Yu, Lee, Huerta-Saenz, and Allen, 2023). Similarly, in Nile tilapia (*Oreochromis niloticus*), our previous work reveals that *sf1* mutation disrupts testicular morphogenesis and downregulates the expression of key steroidogenic enzymes such as *cyp11c1* (Xie et al., 2016). Despite its established central role, the upstream signaling pathways that regulate Sf1 to commit the Leydig lineage remain poorly defined.

Hedgehog (Hh) signaling pathway is an evolutionarily conserved developmental regulator (Y. Zhang and Beachy, 2023). Canonical Hh signaling involves the binding of ligands to Patched (Ptch) receptors, which relieves the suppression on Smoothened (Smo) and ultimately activates Glioma-associated oncogene homolog (Gli) transcription factors to regulate target genes (Briscoe and Thérond, 2013; Finco, LaPensee, Krill, and Hammer, 2015; Ingham, Nakano, and Seger, 2011). In vertebrates, there are three Hh ligands (i.e. Sonic Hh, Indian Hh and Desert Hh [Dhh]), two Ptch receptors (Ptch1 and Ptch2) and three Gli homologs (Gli1-3) (Belgacem and Borodinsky, 2015; Carpenter et al., 1998; Cervantes, Lau, Cano, Borromeo-Austin, and Hebrok, 2010; Kothandapani et al., 2020). A large number of studies have well demonstrated that Dhh signaling pathway is critical in gonadal development (Finco et al., 2015; Umehara et al., 2000). Dhh, secreted by Sertoli cells, is critical for Leydig cell commitment in mammals (Mehta et al., 2021; Pachernegg, Georges, and Ayers, 2022). In mice, mutations of *Dhh* lead to Leydig cell dysfunction, testosterone insufficiency, and infertility (Pierucci-Alves, Clark, and Russell, 2001), while human *DHH* mutations are associated with 46,XY DSD (Wei et al., 2022). Strikingly, these phenotypes mirror those caused by *Sf1* deficiencies, suggesting a potential regulatory nexus between Dhh and Sf1 signaling (Park et al., 2007). However, the molecular circuitry connecting Dhh signaling to Sf1 activation remains elusive.

Studies demonstrate functional divergence between Ptch1 and Ptch2. For instance, *Ptch1* knockout causes embryonic lethality due to constitutive Hh activation, whereas *Ptch2* deficiency yields no overt defects (Cong, Zhu, Chen, Chen, and Chong, 2025; Nieuwenhuis et al., 2006). During testicular development, Ptch1 displays high expression at early stage critical for gonadal primordium formation, while Ptch2 shows testis-enriched dynamic regulation (Kim, Lee, Seppala, Cobourne, and Kim, 2020; H. H. Yao, Whoriskey, and Capel, 2002). However, the distinct roles of Ptch1 and Ptch2 in Leydig lineage commitment remain unknown to date. Among Gli transcription factors, Gli1 acts as a primary activator, while Gli2 and Gli3 exhibit dual regulatory functions (Hui and Angers, 2011; Ruiz i Altaba, Sánchez, and Dahmane, 2002). In murine testes, co-expression of Gli1 and Gli2 in Leydig cells illustrates functional redundancy—single knockouts show normal development, but combined pharmacological inhibition (GANT61) abolishes Leydig cells and spermatogenesis (I. Barsoum and Yao, 2011). These findings underscore the complexity of Gli-mediated transcription in the testis (Kothandapani et al., 2020) and highlight the need to delineate the specific roles of individual Gli in SLC differentiation.

Nile tilapia (*Oreochromis niloticus*) is not only an important farm fish in global aquaculture but also a distinguished animal model for investigating gene function and endocrine regulation. This model organism possesses multiple advantageous attributes, including an XX/XY sex-determination system, early sexual maturity (approximately 3 months after hatching for males and 6 months after hatching for females), a short spawning cycle of approximately 14 days, year-round reproductive capacity under laboratory conditions, well-characterized sex-linked genetic markers, and a body size amenable to physiological sampling (Li, Sun, Zhou, and Wang, 2024; Sun et al., 2014). Furthermore, a stem Leydig cell line (TSL) has been established from the testis of a 3-month-old Nile tilapia (Huang et al., 2020). TSL expresses platelet-derived growth factor receptor α (*pdgfr*α), *nestin*, and chicken ovalbumin upstream promoter transcription factor II (*coup-flla*), which are usually considered as SLC-related markers in several other species (H. Chen, Wang, Ge, and Zirkin, 2017; Ge et al., 2006; Jiang et al., 2014). Notably, this cell line exhibits the capacity to differentiate into 11-ketotestosterone (11-KT)-producing Leydig cells both *in vitro* and *in vivo* (Huang et al., 2020). When cultured in a defined induction medium, TSL cells differentiate into a steroidogenic phenotype, expressing key steroidogenic genes including *star1*, *star2*, and *cyp11c1*, and producing 11-KT; upon transplantation into recipient testes, TSL cells successfully colonize the interstitial compartment, activate the expression of steroidogenic genes, and restore 11-KT production. In this study, by combining CRISPR/Cas9-mediated gene knockout with TSL transplantation technology, we unveil a Dhh-Ptch2-Gli1-Sf1 axis that is essential for Leydig cell lineage differentiation. Our findings provide fundamental insights into the endocrine regulation of Leydig cell development.

## RESULTS

### Mutation of *dhh* disrupts testicular organization and androgen synthesis

The Nile tilapia Dhh open reading frame (GenBank ID: 100707022, 1377 bp) was cloned and validated. The encoded 458-amino acid protein shares over 56% identity with vertebrate homologs (Fig. S1A) and clusters phylogenetically with mammalian DHH orthologs (Fig. S1B). Transcriptomic analyses conducted on various tissues and testes across different developmental stages revealed that *dhh* expression peaked in the testes, particularly at 90 days after hatching (dah) (Fig. S1C, D).

CRISPR/Cas9 targeting of *dhh* exon 1 from Nile tilapia generated homozygous mutants (*dhh*^-/-^) harboring a 13-bp (CAGGGATGCGGAC) frameshift deletion (Fig. S2A–D), resulting in premature termination and loss of the conserved Hedge domain (Fig. S2E). *dhh* mRNA levels in *dhh*^-/-^ fish were significantly reduced compared to WT (Fig. S2F). At 90 dah, *dhh*^-/-^ testes exhibited marked atrophy (Fig. 1A, B) and disorganized histology, with sparse spermatogonia, rare spermatids, and collapsed interstitial architecture (Fig. 1C–D’, K). Immunofluorescence confirmed significantly reduced germ cell (Vasa^+^), meiotic cell (Sycp3^+^), and Leydig cell (Cyp11c1^+^) populations (Fig. 1E-J’, L-N). Serum 11-KT levels in *dhh*^-/-^ fish were significantly lower than WT (Fig. 1O), indicating impaired androgen synthesis.

**Fig. 1.**
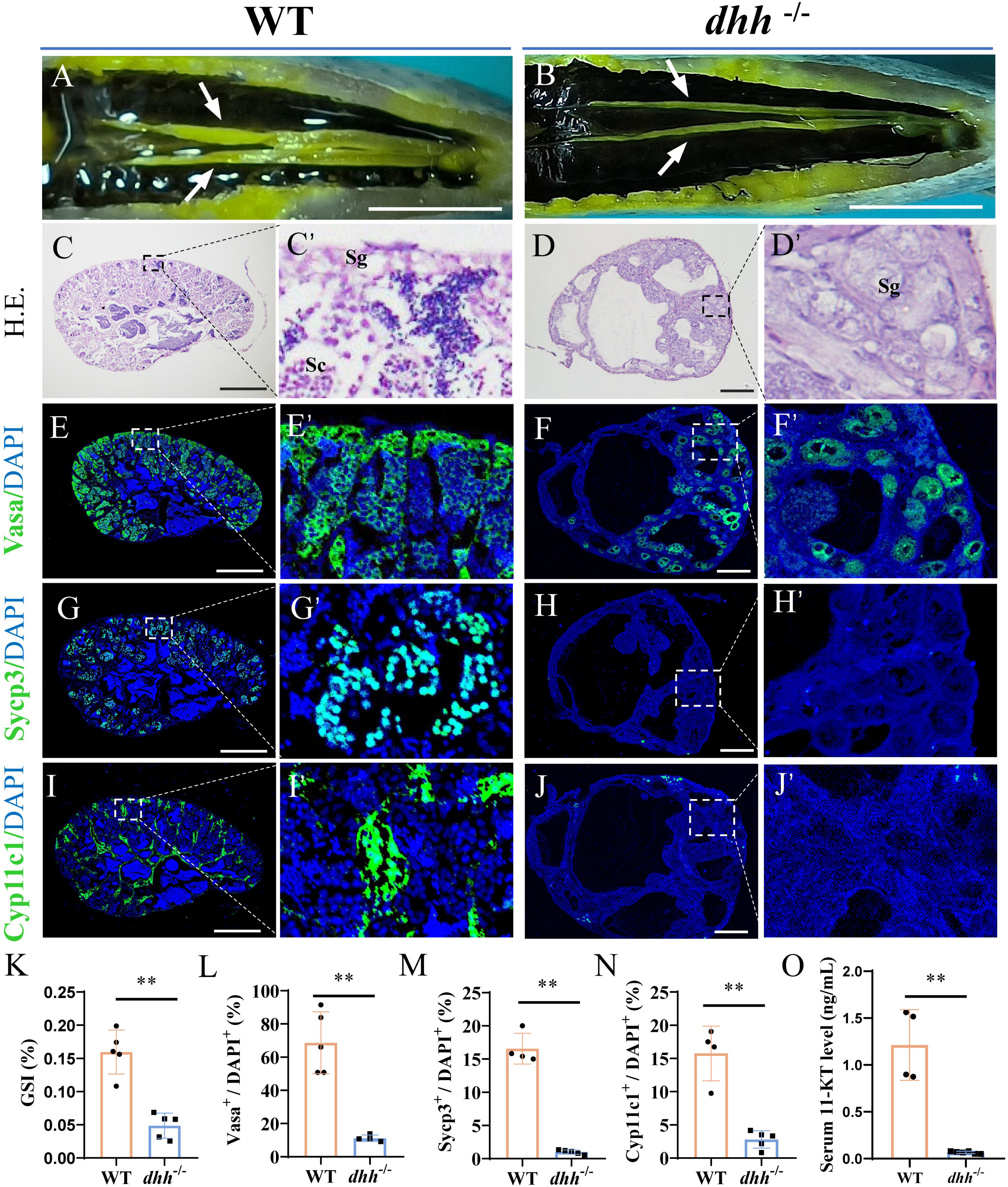
Mutation of *dhh* resulted in testicular developmental disorders. (A-B) Morphological analysis of the testes from WT (*dhh*^+/+^) and *dhh^-^*^/-^ XY fish at 90 dah. Arrows indicate the location of the testes. (C-D’) Histological analysis of testicular sections from WT and *dhh^-^*^/-^ XY fish at 90 dah. Sg, spermatogonia; Sc, spermatocyte. (E-J’) Representative immunofluorescence images showing the expression of germ cell marker Vasa (E-F’), meiosis cell marker Sycp3 (G-H’) and Leydig cell marker Cyp11c1 (I-J’). (K) Gonadosomatic index (GSI) of the testes from WT and *dhh^-^*^/-^ XY fish at 90 dah (*n*=5 fish/genotype). (L-N) Quantification of the percentage of cells positive for Vasa (E-F’), Sycp3 (G-H’) and Cyp11c1 (I-J’) among all DAPI-positive cells in the WT and *dhh^-^*^/-^ testes (*n*=5 fish/genotype). (O) Serum 11-KT level of WT and *dhh^-^*^/-^ XY fish at 90 dah (*n*=6 fish/genotype). Values were presented as mean ± SD. Differences were determined by two-tailed independent Student’s t-test. **, *P* < 0.01. Scale bars, (A-B), 1 cm; (C-J), 50 μm.

### The differentiation of SLCs cannot be rescued by 11-KT, but by SAG

Given the established role of 11-KT in teleost testicular development and spermatogenesis (Zheng et al., 2020), we first asked whether exogenous 11-KT could ameliorate the testicular defects in *dhh*^-/-^ mutants. Compared to untreated mutants, *dhh*^-/-^ individuals exposed to 11-KT exhibited a more organized testicular architecture, with significantly increased populations of Vasa□ and Sycp3□, as evidenced by both H&E staining and immunofluorescence (Fig. 2A–C’, E–G’, I–K’, R, S). This improvement was further supported by a recovery in GSI (Fig. 2Q). However, the population of Cyp11c1-positive Leydig cells remained profoundly depleted in *dhh*^-/-^ +11-KT testes (Fig. 2M–O’, T). Taken together, these data indicate that 11-KT treatment partially restored the development of germ cells but not Leydig cell lineage, implying that the Leydig cell defect is androgen-independent.

**Fig. 2.**
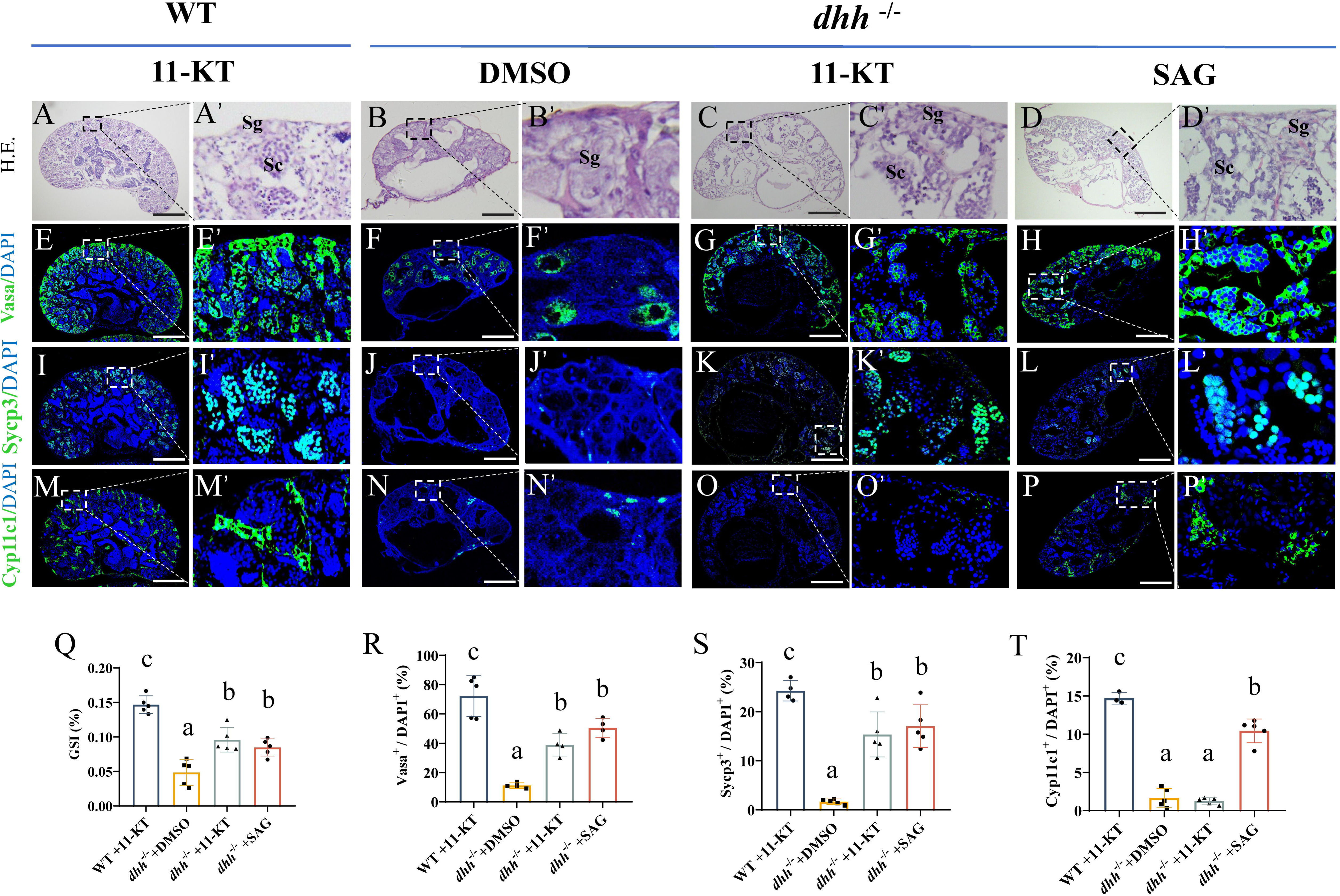
Rescue of testicular development in *dhh^-^*^/-^ XY Nile tilapia by 11-KT and SAG. *dhh^-^*^/-^ XY Nile tilapias at 30 dah were treated either with 11-KT via water immersion (with water changed every two days) or with 10 mg/kg SAG via intraperitoneal injection (with supplemental injection every seven days). WT controls received 11-KT, and *dhh^-^*^/-^ controls received an equivalent volume of the DMSO. Then, at 90 dah, morphological and histological experiments were conducted. (A-D’) Histological analysis of testicular sections from 90 dah XY fish subjected to different treatments as indicated. Sg, spermatogonia; Sc, spermatocyte. Scale bars, 50 μm. (E-P’) Representative immunofluorescence images showing the expression of Vasa (E-H’), Sycp3 (I-L’) and Cyp11c1 (M-P’). Scale bars, 50 μm. (Q) GSI of testes from the different treatment groups at 90 dah (*n*=5 fish per group). (R-T) Quantification of the percentage of cells positive for Vasa (E-H’), Sycp3 (I-L’) and Cyp11c1 (M-P’) among all DAPI-positive cells in the testes (*n*=3~5 fish per group). Values were presented as mean ± SD. Different letters above the error bar indicate statistical differences at *P* < 0.05 as determined by one-way ANOVA followed by Tukey test.

We next investigated whether direct activation of the Dhh pathway could rescue the *dhh*^-/-^ phenotype. Treatment of *dhh*^-/-^ mutants with SAG, a Hedgehog agonist, significantly rescued testicular development. This was marked by a significant recovery of germ cells (Vasa[), spermatocytes (Sycp3[), and critically, Leydig cells (Cyp11c1[), alongside a restoration of testicular histology (Fig. 2D–D’, H–H’, L–L’, P–P’, Q–T). These results demonstrate that *dhh* mutation does not disrupt survival of endogenous Leydig cell lineage, but directly impairs their differentiation, leading to secondary impairment in androgen production and germ cell development.

### Mutation of *dhh* blocks SLCs differentiation *in vivo*

To dissect Leydig cell lineage impairment in *dhh*^-/-^ testes, we transplanted the TSL labeled with PKH26 (a fluorescent red hydrophobic membrane dye that enables tracking of transplanted cells) into WT and *dhh*^-/-^ testes (Fig. 3A). In WT recipients, TSL cells colonized the interstitium and differentiated into Cyp11c1 positive steroidogenic cells (Fig. 3B1–B3). In *dhh*^-/-^ testes, TSL engrafted equivalently but failed to express Cyp11c1 (Fig. 3C1–C3), indicating a differentiation-specific block. Rescue experiments showed that the TSL overexpressing tilapia Dhh (TSL-*On*Dhh) or treated with SAG (TSL+SAG) restored Cyp11c1 expression in *dhh*^-/-^ testes (Fig. 3D1–E3, F) and a significant increase in endogenous 11-KT levels (Fig. 3G). These results suggest that SLC differentiation is inhibited, whereas the survival and engraftment of PKH26-labeled TSL cells were not affected in *dhh*^-/-^ XY tilapia testes. Collectively, these results demonstrate that Dhh signaling is specifically required for the differentiation of SLCs into steroidogenic cells, but is dispensable for their survival or initial recruitment into the testicular niche.

**Fig. 3.**
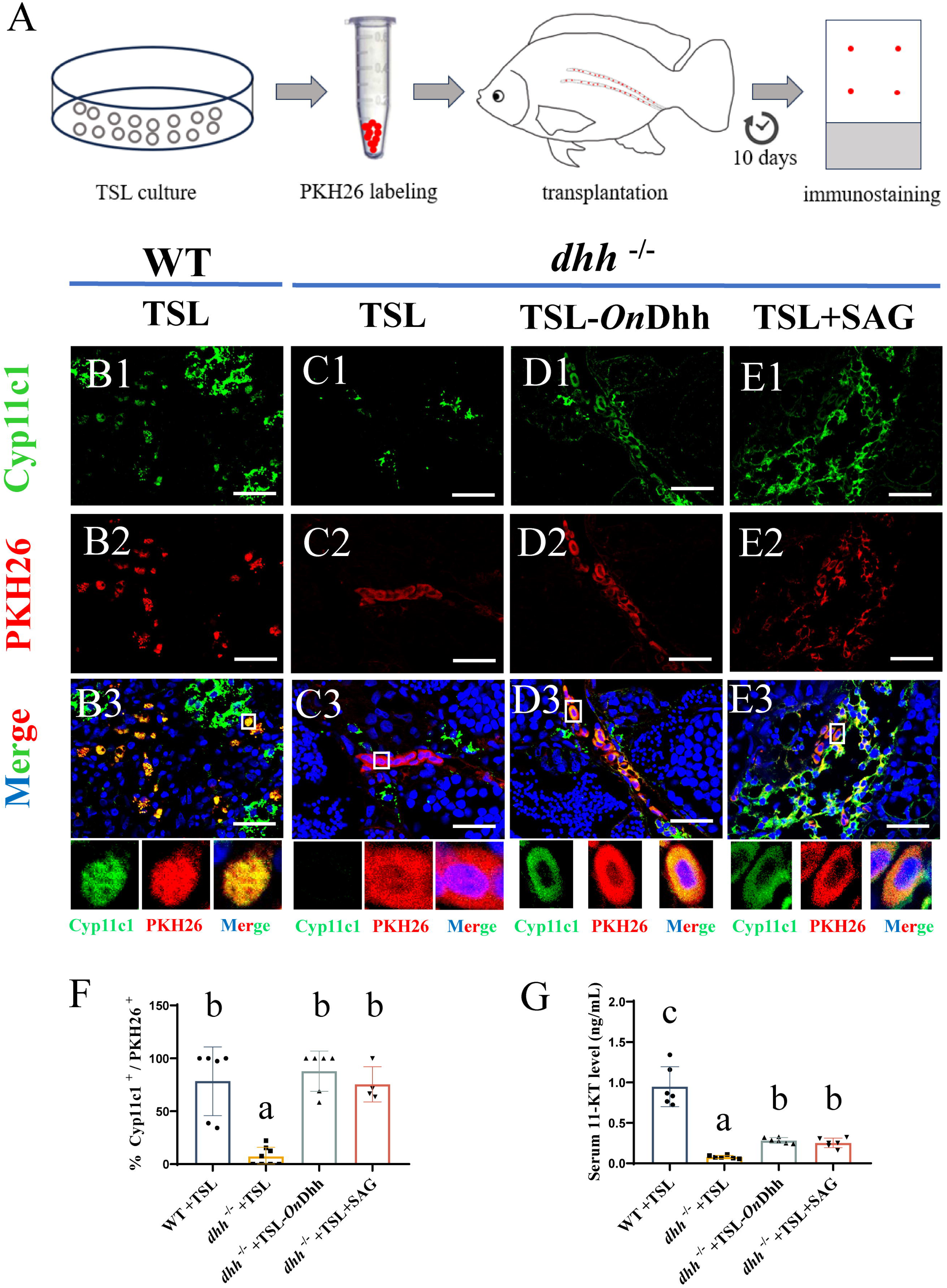
Dhh is required for SLCs differentiation *in vivo*. TSL, Dhh-overexpressing TSL (TSL-*On*Dhh), or 0.5 μM SAG-treated TSL (TSL+SAG) cells were labeled with PKH26 and transplanted into the testes of 90 dah WT or *dhh^-^*^/-^ recipient fish. Analyses were performed 10 days post-transplantation. (A) Schematic diagram of the experimental design for SLCs transplantation and analysis. (B1-E3) Representative immunofluorescence images of testicular sections from recipient fish, showing the localization of transplanted PKH26-labeled SLCs (red) and the expression of Cyp11c1 (green). Nuclei are stained with DAPI (blue). Scale bars, 4 μm. (F) Quantification of the percentage of Cyp11c1-positive cells among the transplanted PKH26-positive SLCs for each treatment group (*n*=5 fish per group). (G) Serum 11-KT level in recipient fish following SLCs transplantation (*n*=5 fish per group). Values were presented as mean ± SD. Different letters above the error bar indicate statistical differences at *P* < 0.05 as determined by one-way ANOVA followed by Tukey test.

### Ptch2 is the functional receptor for Dhh signaling in SLCs

RT-PCR results showed that both *ptch1* and *ptch2* were expressed in TSL (Fig. S3). Fluorescence *in situ* hybridization (FISH) and IF revealed co-expression of *ptch1* and *ptch2* in Cyp11c1 positive Leydig cells (Fig. 4A). To identify the Dhh-driving receptor selectivity in SLCs, we established *ptch1^-/-^* and *ptch2^-/-^* TSL cell lines (Figs. S4A, B) and evaluated pathway activity using an 8×GLI luciferase reporter system (Fig. 4B). In WT TSL cells, the 8×GLI reporter demonstrated substantial luciferase activity that was markedly enhanced by *On*Dhh overexpression (Fig. 4C), confirming Dhh responsiveness in this system. Intriguingly, *ptch1^-/-^* cells maintained Dhh-induced luciferase activation comparable to WT controls, whereas *ptch2^-/-^* cells completely lost this responsiveness (Fig. 4D). This differential receptor requirement implies that Ptch2 likely acts as the functional receptor for transducing Dhh signals in TSL cells.

**Fig. 4.**
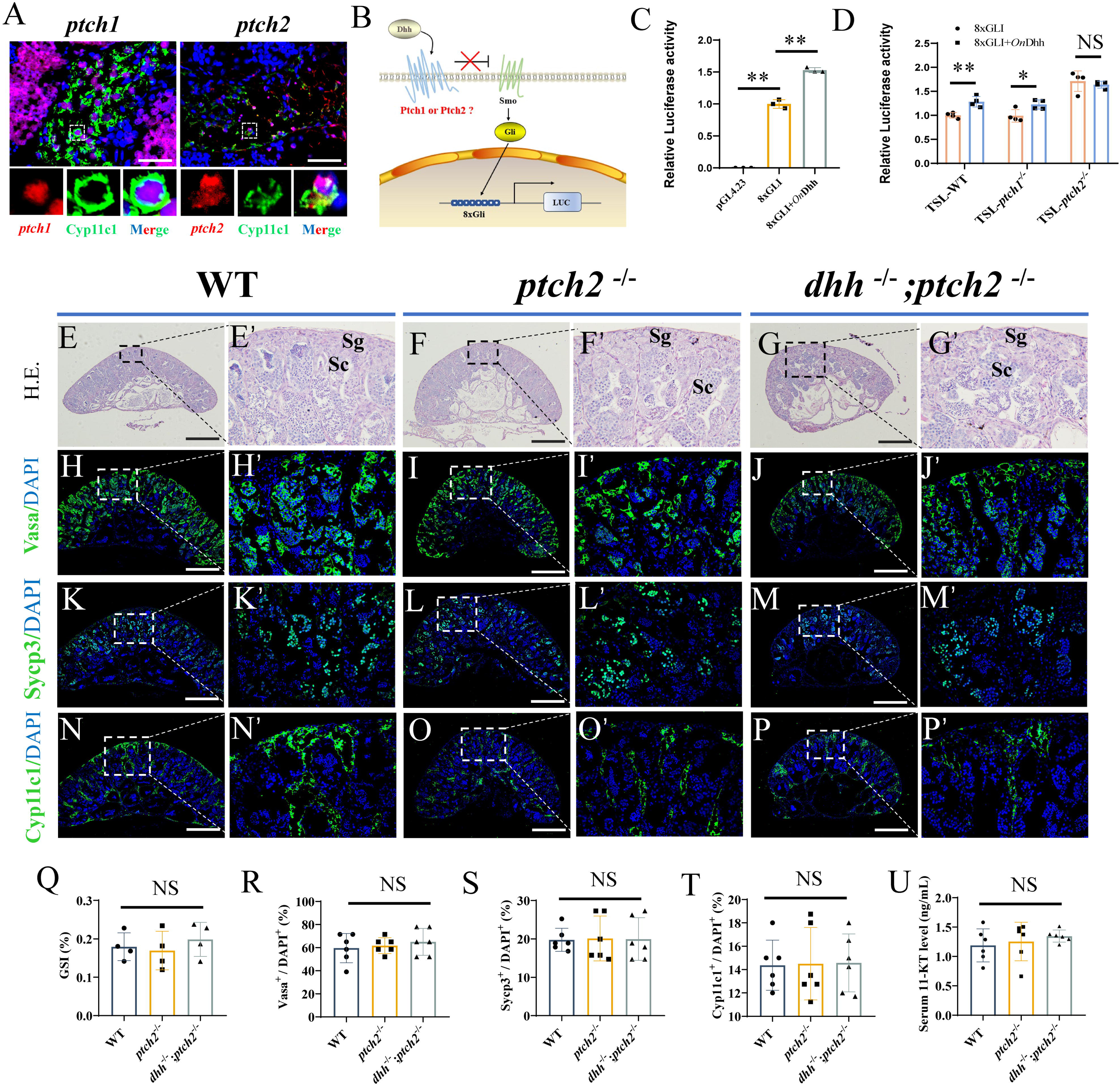
Ptch2 mediates Dhh signaling in SLCs. (A) Co-localization of Cyp11c1 (green, by immunofluorescence) and *ptch1* or *ptch2* mRNA (red, by RNA-FISH) in adult (90 dah) testis sections. Dashed lines outline representative Leydig cells. Scale bars, 4 μm. (B) Schematic illustration of the experimental setup for the luciferase reporter assays shown in panels C and D. (C) Luciferase activity in TSL cells co-transfected with a Gli-responsive reporter (8xGLI) and overexpression of tilapia Dhh (*On*Dhh). A plasmid lacking Gli-binding sites (pGL4.23) served as a negative control (*n*=3). (D) Luciferase activity in TSL-WT, TSL-*ptch1*^-/-^ and TSL-*ptch2*^-/-^ cells transfected with the 8xGLI reporter with or without *On*Dhh (*n*=4). (E-G’) Histological analysis of testis sections from the indicated genotypes at 90 dah. Sg, spermatogonia; Sc, spermatocyte. Scale bars, 50 μm. (H-P’) Immunofluorescence analysis of Vasa (H-J’), Sycp3 (K-M’) and Cyp11c1 (N-P’) in testis sections from the indicated genotypes. Scale bars, 50 μm. (Q) GSI of the testes from the indicated genotypes at 90 dah (*n*=4 fish per group). (R-T) Quantification of the percentage of Vasa (H-J’), Sycp3 (K-M’) and Cyp11c1 (N-P’) positive cells among DAPI-positive cells (*n*=6 fish per group). (U) Serum 11-KT levels in fish of the indicated genotypes at 90 dah (*n*=6 fish per group). Statistical significance was determined by one-way ANOVA followed by Tukey’s test (C, Q-U, different letters above the error bar indicate statistical differences at *P* < 0.05) or Student’s *t*-test (D) (*, *P* < 0.05; **, *P* < 0.01; NS, no significant difference).

To investigate the function of Ptch2 *in vivo*, we generated *ptch2^-/-^* and *dhh^-/-^* ;*ptch2^-/-^* mutant Nile tilapias (Figs. S5A-E). Compared to WT fish at 90 dah, *ptch2^-/-^* testes exhibited no significant differences in testicular histology (Fig. 4E–F’), GSI (Fig. 4Q), germ cell (Vasa^+^, Sycp3^+^) and Leydig cell (Cyp11c1^+^) populations (Fig. 4H-I’, K-L’, N-O’, R-T), and serum 11-KT levels (Fig. 4U). Strikingly, the loss of *ptch2* in the *dhh^-/-^* background (*dhh^-/-^*;*ptch2^-/-^*) fully rescued the testicular defects, resulting in normal histology (Fig. 4G–G’), Leydig cell differentiation (Fig. 4P–P’), and serum 11-KT levels (Fig. 4U). This genetic rescue is consistent with Ptch2 functioning as the inhibitory receptor for Dhh; its removal alleviates the suppression on the pathway, thereby bypassing the requirement for the Dhh ligand.

### Gli1 is the principal transcriptional effector of Dhh signaling in SLCs

In the Hh pathway, Gli transcription factors (Gli1/2/3) mediate diverse developmental processes (26). RT-PCR results showed that *gli1*, *gli2*, and *gli3* were all expressed in TSL (Fig. S3). FISH and IF revealed co-expression of *gli1*, *gli2*, and *gli3* in Cyp11c1 positive Leydig cells (Fig. 5A). To identify Dhh-specific effectors, we generated *gli1^-/-^*, *gli2^-/-^*, and *gli3^-/-^* TSL lines (Fig. S4C–E) and assessed pathway activity using the 8×GLI luciferase reporter. While TSL-*gli2^-/-^* and TSL-*gli3^-/-^*cell lines maintained Dhh-induced luciferase activation comparable to WT controls, the TSL-*gli1^-/-^* cell line completely lost this responsiveness (Fig. 5B), implying that Gli1 is the principal transcriptional effector for Dhh signaling transduction in SLCs.

**Fig. 5.**
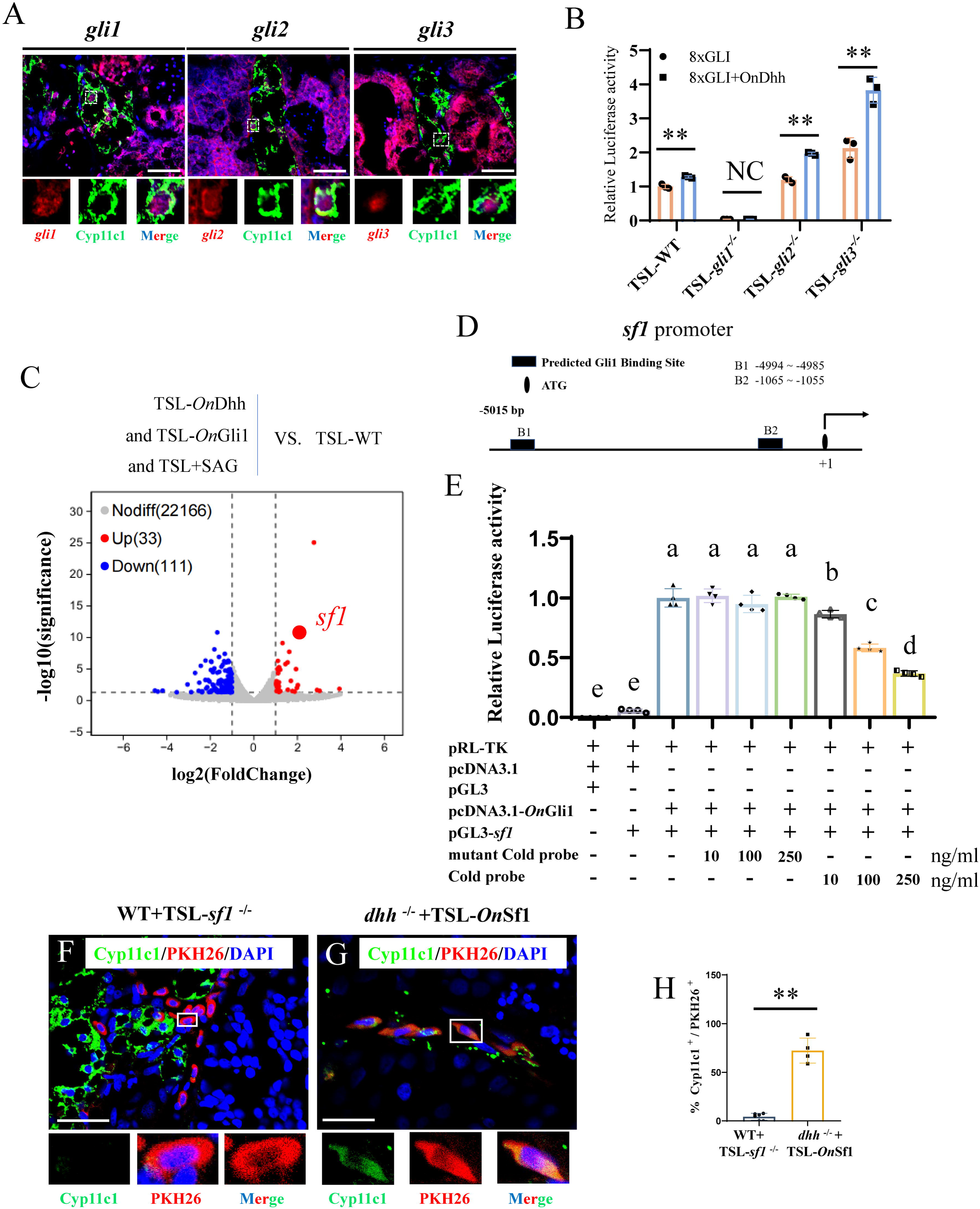
Gli1 transactivates *sf1* to drive SLC differentiation. (A) Co-localization of Cyp11c1 (green) and *gli1*, *gli2*, or *gli3* mRNA (red) in adult (90 dah) testis sections by immunofluorescence and RNA-FISH. Dashed lines outline representative Leydig cells. Scale bars, 4 μm. (B) Luciferase activity in TSL-WT, TSL-*gli1*^-/-^, TSL-*gli2*^-/-^ and TSL-*gli3*^-/-^ cells transfected with the 8xGLI reporter with or without *On*Dhh (*n*=4). (C) Volcano plots of transcriptomic changes in TSL-*On*Dhh, TSL-*On*Gli1 and TSL+ SAG, compared to TSL-WT. Red and blue dots represent significantly up- and down-regulated genes, respectively. (D) Schematic of the tilapia *sf1* gene promoter, indicating the two predicted Gli1 binding sites (B1 and B2). (E) Transcriptional activation of the *sf1* promoter by Gli1. HEK293 cells were co-transfected with pRL-TK, pGL3, pcDNA3.1, pGL3-*sf1*, pcDNA3.1-*On*Gli1, and the indicated cold probe constructs, and luciferase activity was measured 48 hours post-transfection. Competition assays were performed using unlabeled cold probe (Cold probe, GACCACCCA, 10/100/250 ng/mL) or mutant unlabeled cold probe (mutant Cold probe, TTAATTAAA, 10/100/250 ng/mL) (*n*=4). “+” indicates the addition of the corresponding substance, while “-” indicates no addition, and the number represents the amount added (ng/ml). (F-G) Representative immunofluorescence images of testicular sections from recipient fish transplanted with *sf1*-deficient (TSL-*sf1*^-/-^, F) or Sf1-overexpressing (TSL-*On*Sf1, G) SLCs, stained for Cyp11c1 (green) and PKH26 (red). Nuclei are stained with DAPI (blue). Scale bars, 4 μm. (H) Quantification of the percentage of Cyp11c1-positive cells among the transplanted PKH26-positive SLCs in the two transplantation groups (*n*=5 fish per group). Statistical significance was determined by one-way ANOVA followed by Tukey’s test (E, different letters above the error bar indicate statistical differences at *P* < 0.05) or Student’s *t*-test (B, H) (*, *P* < 0.05; **, *P* < 0.01; NS, no significant difference).

### Sf1 is the critical downstream effector of Gli1 in SLCs differentiation

To identify key targets of the Dhh-Gli1 axis, we performed transcriptomic profiling of TSL cells under conditions of pathway activation: Dhh overexpression (TSL-*On*Dhh), Gli1 overexpression (TSL-OnGli1), and SAG treatment (TSL+SAG). Comparative RNA-seq analysis identified a core set of 33 genes consistently upregulated across all three conditions (Fig. 5C, S6A). Among these, *sf1* emerged as a prominently upregulated candidate. Functional annotation of its promoter region identified two conserved Gli1-binding motifs, B1 (AACCACCCA) and B2 (GAGCCACCCA) (Fig. 5D). A dual-luciferase reporter assay confirmed that Gli1 potently transactivates the *sf1* promoter (Fig. 5E). The specificity of this interaction was further demonstrated by a cold probe competition assay: a wild-type oligonucleotide containing the Gli-binding motif (GACCACCCA) competitively inhibited Gli1-driven *sf1* transactivation in a dose-dependent manner, whereas a mutated version (TTAATTAAA) had no effect (Fig. 5E).

The functional necessity of Sf1 was confirmed by transplantation assays using *sf1^-/-^* and Sf1-overexpressing (TSL-*On*Sf1) TSL cells. While TSL-*sf1^-/-^* cells failed to differentiate into Cyp11c1-positive cells even in WT testes, TSL-*On*Sf1 cells successfully differentiated and expressed Cyp11c1 even in the *dhh^-/-^* testicular environment (Fig. 5F–H). These results definitively position Sf1 as the critical downstream effector executing the Dhh-Ptch2-Gli1 signaling cascade during SLCs differentiation.

## DISCUSSION

This study delineates a critical Dhh-Ptch2-Gli1-Sf1 signaling axis essential for SLCs differentiation in Nile tilapia. By combining *dhh*^-/-^ XY models with SLCs transplantation, we definitively demonstrate that Dhh signaling is dispensable for Leydig cell lineage survival but indispensable for its differentiation. Furthermore, our integrated evidence from targeted gene knockout and functional assays suggests that Ptch2 acts as the primary receptor for Dhh signaling in SLCs, and points to Gli1 as the key transcriptional effector mediating the transactivation of *sf1*. Collectively, these findings implicate Sf1 as the critical downstream target bridging Dhh signaling to steroidogenic maturation, thereby resolving key ambiguities in Leydig cell lineage development and providing evolutionary insights into vertebrate reproductive biology.

Consistent with its established role in mammals, our data in Nile tilapia confirm that Dhh function in male testis development is evolutionarily conserved across vertebrates. While mutations in *Dhh* lead to testosterone insufficiency and infertility in mice (Pierucci-Alves et al., 2001) and are associated with 46,XY DSD in humans (Wei et al., 2022), the precise cellular mechanism remained unresolved. Our study demonstrates that *dhh^-/-^*tilapia phenocopy these defects, exhibiting testicular dysgenesis, a profound deficiency of Leydig cells, and drastically reduced 11-KT levels (Fig. 1). To define the onset of Leydig cell differentiation, we performed a developmental time-course analysis. This revealed that Cyp11c1-positive steroidogenic cells first appear in wild-type testes at 30 dah, while being conspicuously absent in *dhh^-/-^* mutants at this same stage (Fig. S7). This clear temporal pattern establishes ∼30 dah as the developmental window when SLCs initiate their differentiation program in the Nile tilapia. The functional indispensability of Dhh at this specific stage was further confirmed by rescue experiments, in which both SAG treatment and transplantation of Dhh-activated TSL cells restored Cyp11c1-positive cell populations in *dhh^-/-^* testes (Fig. 3). Collectively, our findings position Dhh as a crucial niche signal that acts at a defined developmental checkpoint to drive the differentiation of SLCs into steroidogenic Leydig cells.

The functional divergence of Ptch1 and Ptch2 in Hh signaling represents an intriguing aspect of pathway regulation. Structurally, both receptors share conserved 12-transmembrane domains vital for cholesterol transport but diverge in their cytoplasmic C-terminal regions, which are implicated in differential Smoothened inhibition efficiency (Fleet and Hamel, 2018; Qi, Schmiege, Coutavas, and Li, 2018). In mammals, Ptch1 serves as the primary, broad-spectrum regulator of Hh signaling during embryogenesis, whereas Ptch2 often assumes context-dependent roles, such as in primordial germ cell migration via its co-receptor Gas1 (Kim et al., 2020). However, their potential redundancy or specificity within the Leydig cell lineage has remained an open question. Our study provides significant insight into this issue. We confirmed that both *ptch1* and *ptch2* are expressed in Leydig cells (Fig. 4A). Functional interrogation using TSL cell lines revealed a marked difference: genetic ablation of *ptch2* profoundly attenuated Dhh-induced signaling, as measured by a Gli-responsive luciferase reporter, whereas loss of *ptch1* had no detectable effect under the same conditions (Fig. 4B-D). This *in vitro* data, pointing to a primary role for Ptch2, was further supported by *in vivo* genetic interaction studies. The observation that the *dhh^-/-^* phenotype was fully rescued in *dhh^-/-^*;*ptch2^-/-^* double mutants (Fig. 4E-U) is consistent with Ptch2 acting as the inhibitory receptor whose loss alleviates pathway suppression. The specificity for Ptch2 in this context might stem from unique co-receptor interactions or expression patterns within the testicular niche. To preliminarily assess potential compensatory regulation, we examined *ptch1* expression in XY testes from WT, *ptch2*^-/-^ and *dhh^-/-^*;*ptch2^-/-^* fish at 90 dah. No significant differences in *ptch1* mRNA levels were detected among these genotypes (Fig. S8), suggesting that loss of *ptch2* does not trigger compensatory upregulation of *ptch1* at the transcriptional level under the conditions examined. Nonetheless, global *ptch2* mutation affects multiple tissues, whereas our mechanistic focus is on SLC differentiation within the testicular niche. Moreover, the early embryonic lethality of global *ptch1* mutation in tilapia (Liu et al., 2024) precludes direct assessment of its role in postnatal testis development. Therefore, although our findings strongly support a predominant role for Ptch2 in mediating Dhh signaling in SLCs, definitive resolution of receptor specificity will require future Leydig cell-specific conditional knockout models.

Gli is a nuclear transcription factor, characterized by the presence of a zinc finger domain located in the middle of the protein (Kinzler, Ruppert, Bigner, and Vogelstein, 1988). Downstream of the zinc finger domain is a phosphorylation cluster containing phosphorylation sites for protein kinase A and glycogen synthase kinase 3, which negatively regulate the transcriptional activity of Gli through phosphorylation (Niewiadomski et al., 2014). In *Drosophila*, Ci is the sole Gli homolog, possessing both transcriptional activation and repression functions (Huangfu and Anderson, 2006). In vertebrates, there are three Gli homologs: Gli1, Gli2, and Gli3. Gli1 functions solely as a transcriptional activator because it lacks proteolytic processing domains. Gli2 has both activation and repression functions, while Gli3 primarily acts as a transcriptional repressor (Niewiadomski et al., 2019; Pan and Wang, 2007). However, there has been no research on the regulatory mechanism of Gli1/2/3 in SLCs differentiation. In this study, we found that although *gli1*, *gli2*, and *gli3* are all expressed in Leydig cell lineage (Figure 5A-C), the results from TSL cell lines with mutations in *gli1*, *gli2*, and *gli3* indicated that TSL-*gli1^-/-^*, rather than TSL-*gli2^-/-^* or TSL-*gli3^-/-^*, failed to respond to Dhh signaling (Figure 5D). This suggests that Gli1 is the key transcription factor downstream of the Dhh signaling cascade in SLCs.

Sf1 and Dhh play important roles in Leydig cell commitment and testicular development (Parker et al., 2002; Y. Zhang and Beachy, 2023). Studies suggest a potential interaction between these two factors. For example, in *Dhh^-/-^* XY mice, testicular development is impaired, with a significant reduction in the number of Leydig cells and a marked downregulation of *Sf1* expression (H. H. Yao et al., 2002). In XX mouse ovaries where Smo is constitutively activated, a large number of ectopic Sf1-positive Leydig cells are generated (I. B. Barsoum, Bingham, Parker, Jorgensen, and Yao, 2009). Similarly, in *sf1^-/-^* XY tilapia, testicular development is impaired, with a significant reduction in Leydig cell numbers and down regulation of *dhh* expression (Xie et al., 2016). These findings imply that Dhh and Sf1 may interact through feedback loops or co-regulatory mechanisms. However, direct evidence for the regulation of Sf1 by the Hh signaling pathway remains lacking. This study elucidates the molecular mechanism of Sf1 as a critical downstream effect of the Dhh-Ptch2-Gli1 axis. Transcriptomic profiling revealed an increased expression of *sf1* by Dhh regulation signaling in TSL cells (Fig. S6), identifying it as a key downstream target of Dhh signaling. Promoter analysis identified two conserved Gli-binding motifs (Fig. 5F) within the *sf1* promoter, and luciferase assays confirmed that Gli1 transactivates *sf1* transcription (Fig. 5G). Functional validation demonstrated that TSL-*sf1^-/-^* cells failed to differentiate even in WT testes, whereas TSL-OnSf1 cells restored Leydig cell differentiation and steroidogenesis in *dhh^-/-^* testes (Fig. 5H-J). These results establish Sf1 as both necessary and sufficient for Leydig cell lineage differentiation.

In conclusion, our study systematically deciphers the signaling cascade governing SLCs differentiation, from the niche signal to the core steroidogenic regulator. We definitively show that the Dhh signal from the niche is indispensable for SLCs differentiation, but not their survival. Within this pathway, Ptch2 serves as the critical receptor, relaying the signal through the transcription factor Gli1. The pivotal outcome of this cascade is the activation of *sf1* by Gli1. Thus, we establish a complete Dhh-Ptch2-Gli1-Sf1 axis, resolving the long-standing question of how a key morphogen controls Leydig cell lineage development in vertebrates.

## MATERIALS AND METHODS

### Animals

Nile tilapias (*Oreochromis niloticus*) were kept in recirculating freshwater systems at 26□ under a 12-h light/dark photoperiod. All animal experiments were conducted in accordance with the regulations of the Guide for Care and Use of Laboratory Animals prescribed by the Committee of Laboratory Animal Experimentation at Southwest University, China (IACUC-20181015-12).

### Generation of mutant lines by CRISPR/Cas9 in Nile tilapia

The sequences of tilapia *dhh* (GenBank ID: 100707022) and *ptch2* (GenBank ID: 100692939) were retrieved from the NCBI. CRISPR/Cas9-mediated mutagenesis was performed as described previously (Li et al., 2014). Guide RNAs (gRNAs) targeting the first exon of *dhh* (GCGGGCCCGGTCCGCATCCC) and *ptch2* (GTCCCAGGGGCCGGCGTATT) were designed using ZiFit (http://zifit.partners.org/ZiFiT/). gRNA and Cas9 mRNA were synthesized following established protocols (Li et al., 2014). The primers listed in Table S1. A mixture of synthetic gRNA (500 ng/μl) and Cas9 mRNA (1000 ng/μl) was microinjected into one-cell-stage fertilized embryos. Mutants were validated by polyacrylamide gel electrophoresis (PAGE) and Sanger sequencing. F0 chimeric XY males were crossed with WT XX females to generate heterozygous F1 offspring. F1 siblings carrying a 13-bp deletion in *dhh* or a 25-bp deletion in *ptch2* were intercrossed to generate homozygous F2 mutants. For double mutants, *dhh*^+/-^ XY males were mated with *ptch2*^+/-^ XX females to obtain *dhh*^+/-^;*ptch2*^+/-^ offspring. These double heterozygotes were then intercrossed to generate *dhh*^-/-^*;ptch2*^-/-^ mutants.

### Generation of mutant lines by pCas9-NtU6sgRNA system in TSL Cells

The tilapia stem Leydig cell line TSL was maintained in ESM4 medium as described (Huang et al., 2020). Gene sequences for *ptch1* (GenBank ID: 100700194), *ptch2* (GenBank ID: 100692939), *gli1* (GenBank ID: 100710687), *gli2* (GenBank ID: 100704788), *gli3* (GenBank ID: 100707331), and *sf1* (GenBank ID: 100534561) were retrieved from NCBI. Knockout plasmids targeting these genes were designed following established protocols (Z. Zhang et al., 2023). gRNAs were generated using ZiFit, with the following exon-specific targets: *ptch1* (GGAGGCGCTCCTGCAGCACC, exon 3), *ptch2* (GTCCCAGGGGCCGGCGTATT, exon 1), *gli1* (TGACTCATGGATCAGGACCA, exon 2), *gli2* (GATTCGCCTGTCACCCCACG, exon 3), *gli3* (TGAGGAGCCCTCTACGTCTA, exon 2), and *sf1* (GGCTGTGTACTGGTACTGGG, exon 4; designed as previously described) (Z. Zhang et al., 2023). Primers are listed in Table S1. Plasmids were transfected into TSL cells using the MDMP E-1 microchannel transfection instrument (MENDGENE) with transwell cell compartments (BIOFIT), per manufacturer guidelines. Post-transfection (48 hours), cells were selected in ESM4 medium containing 500 μg/mL G418 for 7 days. Single-cell clones were isolated via dilution and serial passage, followed by genomic DNA (gDNA) extraction and Sanger sequencing to confirm mutations.

### Histological analyses of testes from WT, *dhh^-/-^*, *ptch2^-/-^* and *dhh^-/-^;ptch2^-/-^* XY fish

GSI was calculated for WT, *dhh^-/-^*, *ptch2^-/-^* and *dhh^-/-^; ptch2^-/-^* XY fish at 90 dah as: (gonad weight / body weight) × 100%. Testes were fixed in Bouin’s solution for 24 hours at room temperature with agitation, dehydrated through an ethanol series, and embedded in paraffin. Sections (5 μm thickness) were stained with hematoxylin and eosin (H&E) and imaged using an Olympus BX53 microscope. Germ cell classification (spermatogonia, spermatocytes) followed established criteria (Zhao et al., 2023).

### Immunofluorescence

Testes from WT, *dhh^-/-^*, WT+11-KT, *dhh^-/-^*+11-KT, *ptch2^-/-^* and *dhh^-/-^; ptch2^-/-^* XY fish at 90 dah were analyzed for germ cell marker Vasa, Leydig cell marker Cyp11c1, and meiosis marker Sycp3. Tissue fixation and embedding. Sections (5 μm) were deparaffinized, rehydrated, and subjected to antigen retrieval. After blocking with 5% bovine serum albumin (BSA; 30 minutes, room temperature), sections were incubated overnight at 4°C with rabbit polyclonal primary antibodies (Vasa, Cyp11c1, Sycp3; 1:1,000 dilution), validated previously (Dai et al., 2021). Alexa Fluor 488-conjugated goat anti-rabbit secondary antibody (1:1,000; Invitrogen) was applied for 1 hour at 37°C, followed by DAPI nuclear counterstaining (1:1,000; Sigma-Aldrich). Fluorescence signals were imaged using an Olympus Fv3000 confocal microscope. For each biological replicate (*n*=5-6 fish per genotype), three non-serial, non-adjacent testis sections were analyzed. From each section, three representative fields of view were captured to ensure non-overlapping sampling. All positive cells number of Vasa, Sycp3 and Cyp11c1 was quantified by Image J Pro 1.51 software using default parameters.

### Fluorescence *in situ* hybridization (FISH) and immunofluorescence

Testes from WT XY tilapia at 90 dah were fixed in 4% paraformaldehyde (Sigma) overnight at 4°C with agitation, paraffin-embedded, and sectioned (5 μm). FISH was performed as described (Zhao et al., 2023). Partial cDNA fragments of *ptch1*, *ptch2*, *gli1*, *gli2*, and *gli3* were cloned into the pGEM-T Easy vector. Sense (control) and antisense RNA probes were synthesized using a DIG RNA Labeling Kit (Roche) and T7 mMESSAGE mMACHINE Kit (Ambion). Signals were amplified with the TSA^TM^ Plus TMR system (PerkinElmer) per manufacturer protocols (primers in Table S1). Concurrently, Leydig cell marker Cyp11c1 expression was assessed by immunofluorescence.

### 11-KT and SAG rescue experiments

To rescue *dhh^-/-^* XY fish, offspring from *dhh*^+/-^ XY male × *dhh*^+/-^ XX female crosses were divided into two groups. The experimental group received 11-KT via immersion from 30 to 90 dah, with treatments refreshed every 2 days. 11-KT concentrations increased incrementally from 57 ng/L at 30 dah to 1254 ng/L by 90 dah (39.9 ng/L increase per treatment cycle; see Fig. 2, 5, S7). For SAG rescue, fish received intraperitoneal injections of 10 mg/kg SAG from 30 to 90 dah, with booster doses administered weekly. Testes were collected at 90 dah for morphological and histological evaluation.

### TSL cell labeling, transplantation and IF

TSL, previously isolated from 3-month-old XY tilapia testes (Huang et al., 2020), was used to assess stem Leydig cell differentiation in WT and *dhh^-/-^* XY fish. TSL cells were treated as follows: (1) untreated TSL, (2) TSL + 0.5 μM SAG (Hh pathway agonist), (3) TSL-*On*Dhh (overexpressing tilapia Dhh), (4) TSL-*On*Sf1 (overexpressing tilapia Sf1), and (5) TSL-*sf1^-/-^* (*sf1* mutation). For the SAG treatment experiment, TSL cells were incubated with 0.5 μM SAG for 48 hours before transplantation. Cells were labeled with PKH26 fluorescent dye (Sigma) and transplanted (5,000 cells per testis) into WT or *dhh^-/-^* XY tilapia via the urogenital tract (Huang et al., 2020). For TSL-*On*Dhh, the Dhh ORF (1377 bp) was amplified from gDNA, cloned into pcDNA3.1 (Invitrogen), and transfected into TSL cells (primers in Table S1). At 10 days post-transplantation, testes were analyzed by IF to quantify Cyp11c1 positive cells among PKH26 positive populations.

### GLI report assay

A firefly luciferase reporter plasmid (8×GLI) was generated by cloning eight tandem copies of the GLI-binding motif (GACCACCCA) into the pGL4.23 vector (minimal promoter). To validate 8×GLI functionality, TSL cells were transiently transfected with: (1) pGL4.23 (negative control), (2) 8×GLI, or (3) 8×GLI plus Dhh-overexpression plasmid (*On*Dhh), for 48 hours. The 8×GLI construct was further transfected into TSL (WT), TSL-*ptch1^-/-^*, TSL-*ptch2^-/-^*, TSL-*gli1^-/-^*, TSL-*gli2^-/-^* or TSL-*gli3^-/-^*, with or without *On*Dhh. Transfections included pRL-TK as an internal control. Luciferase activity was measured 48 hours post-transfection using the Dual Luciferase Assay System (Promega) and a Luminoskan™ Ascent luminometer (Thermo Fisher Scientific). Relative activity was calculated as firefly/Renilla luciferase ratios.

### Western blot

The ORFs of Dhh (1377 bp) and Gli1 (4404 bp) were amplified from gDNA by PCR. Two tandem FLAG tag sequences (GATTATAAAGATGATGATAAA) were inserted flanking each ORF (after the start codon and before the stop codon) and cloned into the pcDNA3.1 vector (Invitrogen), generating pcDNA3.1-*On*Dhh and pcDNA3.1-*On*Gli1 (primers in Table S1). TSL cells were transiently transfected with pcDNA3.1-*On*Dhh or pcDNA3.1-*On*Gli1 for 48 hours. Protein expression was analyzed by Western blot following established methods (Dai et al., 2021). Briefly, total protein lysates were separated by 15% SDS-PAGE, transferred to nitrocellulose membranes, and blocked with 5% BSA in TBST (10 mM Tris pH 7.9, 150 mM NaCl, 0.1% Tween-20) for 1 hour at 37°C. Membranes were incubated overnight at 4°C with rabbit anti-FLAG antibody (1:1000; Cell Signaling Technology), followed by HRP-conjugated goat anti-rabbit secondary antibody (1:1000; Invitrogen) for 1 hour at 37°C. Signals were detected using the BeyoECL Plus Kit (Beyotime) and imaged on a Fusion FX7 system (Vilber Lourmat).

### Measurement of 11-KT levels by ELISA

Blood was collected from the caudal vein of 90 dah XY fish (WT, *dhh^-/-^*, *ptch2^-/-^*, *dhh;ptch2^-/-^*, WT+TSL, *dhh^-/-^*+TSL, *dhh^-/-^*+TSL+SAG, *dhh^-/-^*+TSL-*On*Dhh and *dhh^-/-^* +TSL-*sf1^-/-^*), incubated at 4°C overnight, and centrifuged to isolate serum. Tissue fluid samples from WT and *dhh^-/-^* XY fish at 5, 10, 20, and 30 dah were similarly prepared. All samples were stored at −80°C until analysis. Serum and tissue fluid 11-KT levels were quantified using a 11-keto Testosterone ELISA Kit (Cayman) validated for teleost samples, per manufacturer instructions. Individual samples (serum or tissue fluid; *n* = 6 per genotype) were analyzed in triplicate on a single 96-well plate, with concentrations calculated from standard curves.

### Transcriptome analyses

Tissue- and stage-specific expression of *dhh* was assessed using transcriptomic data from tilapia tissues (brain, heart, head kidney, kidney, liver, muscle, ovary, testis) and gonads at developmental stages spanning 5 to 360 dah (Tao et al., 2018; Tao et al., 2013). For global gene expression profiling, RNA was extracted from TSL-WT, TSL-*On*Dhh, TSL-*On*Gli1, and TSL + 0.5 μM SAG cells using RNAiso Plus (Takara). For the SAG treatment experiment, TSL cells were incubated with 0.5 μM SAG for 48 hours before collection. For each genotype, cells from three independent culture wells were pooled. Libraries were sequenced on an Illumina platform (GENEBOOK Biotechnology), and clean reads were aligned to the reference genome with HISAT2. Gene expression levels (FPKM: fragments per kilobase per million mapped reads) were quantified using featureCounts. Differentially expressed genes (DEGs) were identified for each condition (TSL-*On*Dhh, TSL-*On*Gli1, TSL+SAG) compared to TSL-WT controls using edgeR (threshold: FDR < 0.05, |log2(foldchange)| ≥ 1.5).

### Real-time PCR

Testes from WT and *dhh^-/-^* XY fish at 90 dah (*n* = 3 per genotype), along with TSL-WT, TSL-*On*Dhh, TSL-*On*Gli1, and TSL + 0.5 μM SAG cells, were analyzed for gene expression. Total RNA was extracted using RNAiso Plus (Takara), reverse-transcribed into cDNA (PrimeScript II 1st Strand cDNA Synthesis Kit; Takara), and amplified by real-time PCR on an ABI-7500 system (Applied Biosystems). Relative mRNA levels were normalized to β*-actin* and calculated via the formula R = 2^−ΔΔCt^ (primers in Table S1).

### Dual luciferase report assay

The 5015-bp promoter region upstream of the *sf1* start codon was amplified and cloned into the pGL3-basic vector (Promega), generating pGL3-*sf1* (primers in Table S1). To assess Gli1 binding to the *sf1* promoter, HEK293 cells were co-transfected with pGL3-*sf1*, pcDNA3.1- *On*Gli1, cold probe (GACCACCCA), and mutant cold probe (TTAATTAAA) using the TransIT-X2 transfection reagent (Mirus Bio). Control transfections included empty pcDNA3.1 and pGL3 vectors, with pRL-TK for normalization. Luciferase activity was measured 48 hours post-transfection.

### Statistical analyses

Data are presented as mean ± SD from at least three independent replicates. A two-tailed independent Student’s t-test was used to determine the differences between the two groups. One-way ANOVA, followed by Tukey multiple comparison, was used to determine the significance of differences in more than two groups. *P* < 0.05 was used as a threshold for statistically significant differences.

## Supporting information

Graphical abstract

## Data availability

All data generated or analyzed during this study are included in the manuscript and its supporting files. Materials generated in this study are available from the corresponding author upon reasonable request.

## Acknowledgements

This research was funded by the National Key Research and Development Program of China (Grant No. 2022YFD1201600), the National Natural Science Foundation of China (Grant No. 32172969, 31972776, 32102780, 32473159).

## Author contributions

**Changle Zhao**: Writing – original draft, Software, Resources, Methodology, Investigation, Formal analysis, Data curation. **Yongxun Chen**: Writing – original draft, Methodology, Data curation. **Lei Liu**: Data curation, Conceptualization. **Xiang Liu**: Software, Investigation. **Hesheng Xiao**: Software, Data curation. **Feilong Wang**: Investigation, Formal analysis. **Qin Huang**: Resources, Software. **Xiangyan Dai**: Resources, Investigation. **Wenjing Tao**: Resources, Funding acquisition. **Deshou Wang**: Supervision, Investigation, Funding acquisition. **Jing Wei**: Writing – review & editing, Validation, Supervision, Project administration, Funding acquisition, Data curation, Conceptualization.

## Figure legends

**Fig. S1. Sequence analysis and mRNA expression profile of Nile tilapia *dhh*.**

(A) Dhh amino acid sequence alignment. Amino acids are numbered in the right margin. Deletions are indicated by dashes, shaded areas indicate shared sequences. The amino-terminal hedge domain is underlined by dotted line, with the signal peptide region in the front and the autocatalytic carboxy-terminal domain (hog) in the back. The box indicates the autocatalytic site of an absolutely conserved Gly-Cys-Phe tripeptide. At the end of the alignment are percentage identity values of the full-length and hedge domain of tilapia Dhh to the orthologue from other species. (B) Phylogenetic analysis of Dhh. The phylogenetic tree was constructed using the neighbor-joining method within the MEGA7.0 program. Node values represent percent bootstrap confidence derived from 1000 replicates. (C-D) The RPKM (reads per kb per million reads) values of tilapia *dhh* in various adult tissues (C) and XY and XX gonads at 5, 7, 20, 30, 90, 180, 300 dah (days after hatching) (D) in transcriptome sequencing data which were sequenced using Illumina 2000 HiSeq technology in our previous study (Tao et al., 2013).

**Fig. S2. Establishment of the Nile tilapia *dhh* mutant line by CRISPR/Cas9.**

(A) Schematic representation of gRNA targeting the Nile tilapia *dhh* locus. The gRNA was designed to target exon 1. The translation start codon ATG and stop codon TAA were indicated by arrows. The PAM (protospacer adjacent motif) site was marked by green box. DNA sequence alignment of the three mutant lines with WT. The added sequences were marked by red letter, and the deleted sequences were indicated in red dotted lines. One had a 4-bp addition, one had an 8-bp deletion, and the other had a 13-bp deletion. (B) Schematic diagram showing the breeding plans of *dhh* F0 to F2 fish. (C) Homozygous mutants of F2 fish were screened by PAGE. The first lane is DNA marker, the second lane is *dhh^+/+^* fish, the third lane is *dhh^+/-^* fish, and the fourth lane is *dhh^-/-^* fish. (D) Sanger sequencing results of *dhh* genes from WT and the 13-bp deletion homozygous mutant fishes. (E) The *dhh* ORF sequence in WT was indicated in black and the frameshift-altered sequence for the 13-bp deletion *dhh* ORF was indicated in red. (F) The relative mRNA expression of *dhh* in WT and the 13-bp deletion homozygous mutant fishes by RT-qPCR. Values were presented as mean ± SD (*n*=3). Differences were determined by two-tailed independent Student’s t-test. **, *P* < 0.01.

**Fig. S3. RT-PCR analyses of Hh pathway genes in TSL cells.**

β*-actin* served as a loading control, and RNAs from TSL cells were used as templates for negative controls. The numbers in parentheses indicate the PCR cycles.

**Fig. S4. Establishment of the TSL *ptch1*, *ptch2*, *gli1*, *gli2*, *gli3* and *sf1* mutant lines.**

(A-F) Schematic representation of gRNAs targeting the *ptch1*, *ptch2*, *gli1*, *gli2*, *gli3* and *sf1* locus. The gRNAs were designed to target ORF (open reading frame). The PAM (protospacer adjacent motif) sites were marked by red letter. Sanger sequencing of *ptch1*, *ptch2*, *gli1*, *gli2*, *gli3* and *sf1* single mutant alleles in single cell clones. PCR amplicons from DNA templates of the six cell clones were directly used for sequencing.

**Fig. S5. Establishment of the Nile tilapia *ptch2* mutant line by CRISPR/Cas9 system.**

(A) Schematic representation of gRNA targeting the Nile tilapia *ptch2* locus. The gRNA was designed to target exon 1. The translation start codon ATG and stop codon TGA were indicated by arrows. The PAM site was marked by green box. DNA sequence alignment of the five mutant lines with WT. The added sequences were marked by red letter, and the deleted sequences were indicated in red dotted lines. (B) Schematic diagram showing the breeding plans of *ptch2* F0 to F2 fish. (C) DNA sequencing showed the 25-bp deletion within the *ptch2* ORF in the homozygous mutant compared to the WT. (D) Schematic of the prediction of the intact Ptch2 protein in the WT and the truncated Ptch2 protein in the homozygous mutants. (E) Homozygous mutants of F2 fish were screened by PAGE. The first lane is DNA marker, the second lane is *ptch2^+/+^* fish, the third lane is *ptch2^+/-^* fish, and the fourth lane is *ptch2^-/-^* fish. “*” indicates heteroduplex, and arrows indicate homoduplex.

**Fig. S6. Transcriptome data analyses and verification by RT-qPCR.**

(A) Hierarchical clustering analysis of global gene expression patterns in TSL-WT, TSL-*On*Dhh, TSL-*On*Gli1 and TSL+ 0.5 μM SAG. Blue indicates decreased expression, and red indicates increased expression. (B) Heatmap showing the expression patterns of upregulated differentially expressed genes (DEGs) identified in Fig. 5C. The FPKM value for each gene in each sample is indicated within the squares. The color gradient from blue to red reflects low to high expression levels per row (gene). (C) qRT-PCR validation. Values were presented as mean ± SD (n = 3).

**Fig. S7. The expression levels of Cyp11c1 in the testis and the levels of 11-KT in tissue fluid from WT and *dhh*^-/-^ XY fish at 5, 10, 20, and 30 dah.**

(A-E) Representative immunofluorescence images showing Cyp11c1 expression (green) in testis sections from WT fish at 5, 10, 20, 30 dah and *dhh*^-/-^ mutants at 30 dah. Scale bars, 4 μm. (F) Tissue fluid 11-KT levels in WT and *dhh*^-/-^ XY fish at indicated time points (*n*=6 fish per group). Values were presented as mean ± SD. Differences were determined by two-tailed independent Student’s t-test. NS, no significant difference.

**Fig. S8. Expression analysis of *ptch1* in WT and mutant XY testes.**

Relative mRNA expression levels of *ptch1* in XY testes from WT, *ptch2*^-/-^ and *dhh^-/-^*;*ptch2^-/-^* fish at 90 dah by RT-qPCR. Values were presented as mean ± SD (*n*=3 fish per group). Statistical significance was determined by one-way ANOVA followed by Tukey’s test. NS, no significant difference.

**Fig. S9. Validation of key gene expression changes in mutant and overexpression TSL cell lines. (only for reviewer).**

(A) Relative mRNA expression levels of *sf1* in TSL-WT, TSL-*gli1*^-/-^, TSL-*gli2*^-/-^ and TSL-*gli3*^-/-^ cells by RT-qPCR. Values were presented as mean ± SD (*n*=3). Different letters above the error bar indicate statistical differences at *P* < 0.05 as determined by one-way ANOVA followed by Tukey test. (B) Relative mRNA expression of *dhh* in TSL-WT and TSL-*On*Dhh cells by RT-qPCR. Values were presented as mean ± SD (*n*=3). Differences were determined by two-tailed independent Student’s t-test. **, *P* < 0.01. (C) Relative mRNA expression of *sf1* in TSL-WT and TSL-*On*Sf1 cells by RT-qPCR. Values were presented as mean ± SD (*n*=3). Differences were determined by two-tailed independent Student’s t-test. **, *P* < 0.01.

**Fig. S10. Negative control experiments for RNA fluorescence in situ hybridization (FISH). (only for reviewer)**

Co-staining of Cyp11c1 (by immunofluorescence) and sense RNA probes for *ptch1*, *ptch2*, *gli1*, *gli2*, or *gli3* (by FISH) on testis sections from 90 dah fish. The sense probes serve as negative controls to confirm the specificity of the antisense probe hybridization signals shown in the main figures. Scale bars, 4 μm.

