## Supplementary figures and images for "Desert Hedgehog mediates stem Leydig cell differentiation through Ptch2/Gli1/Sf1 signaling axis"

### Graphical abstract

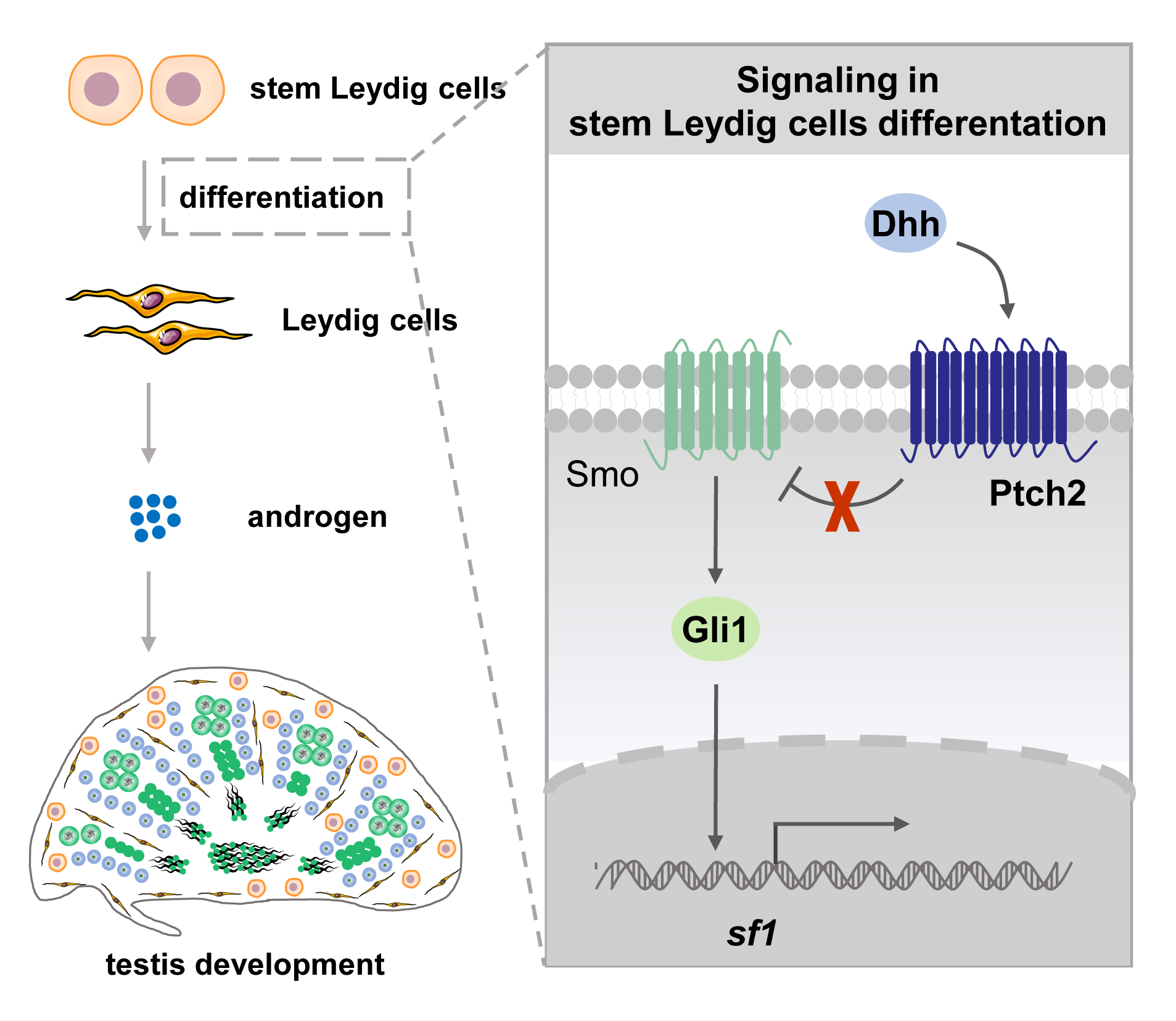

### Supplemental Figure S1

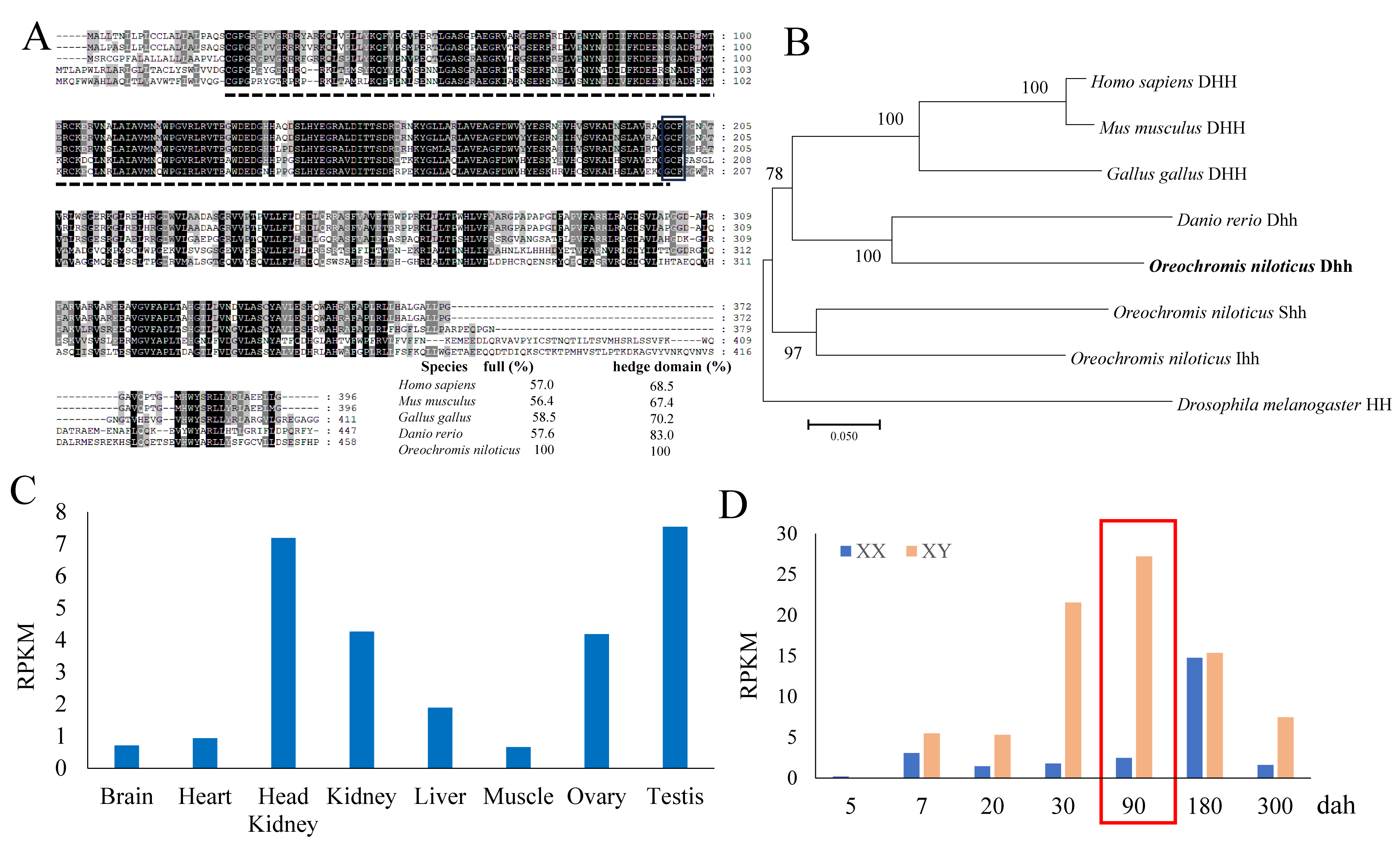

### Supplemental Figure S2

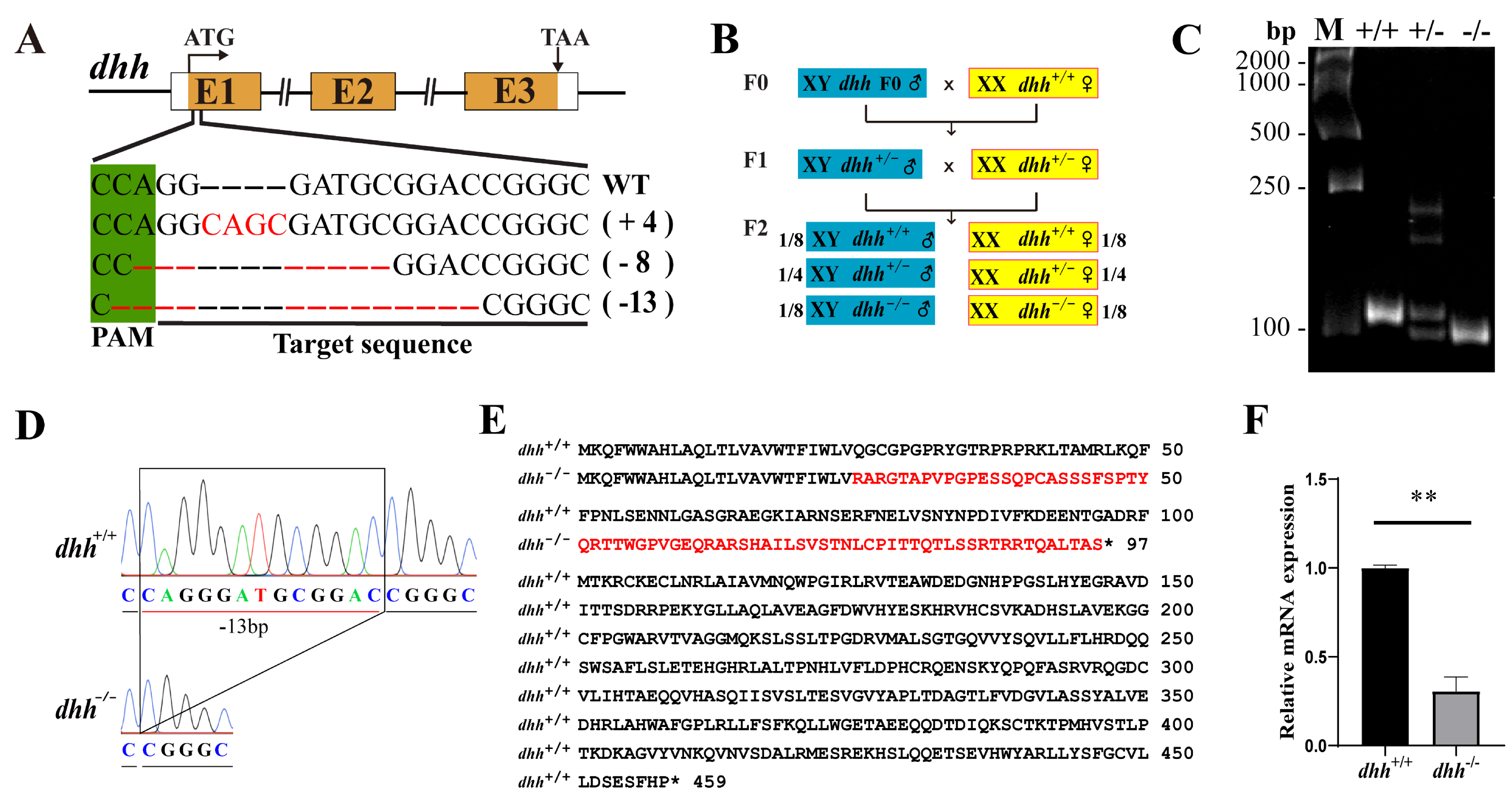

### Supplemental Figure S3

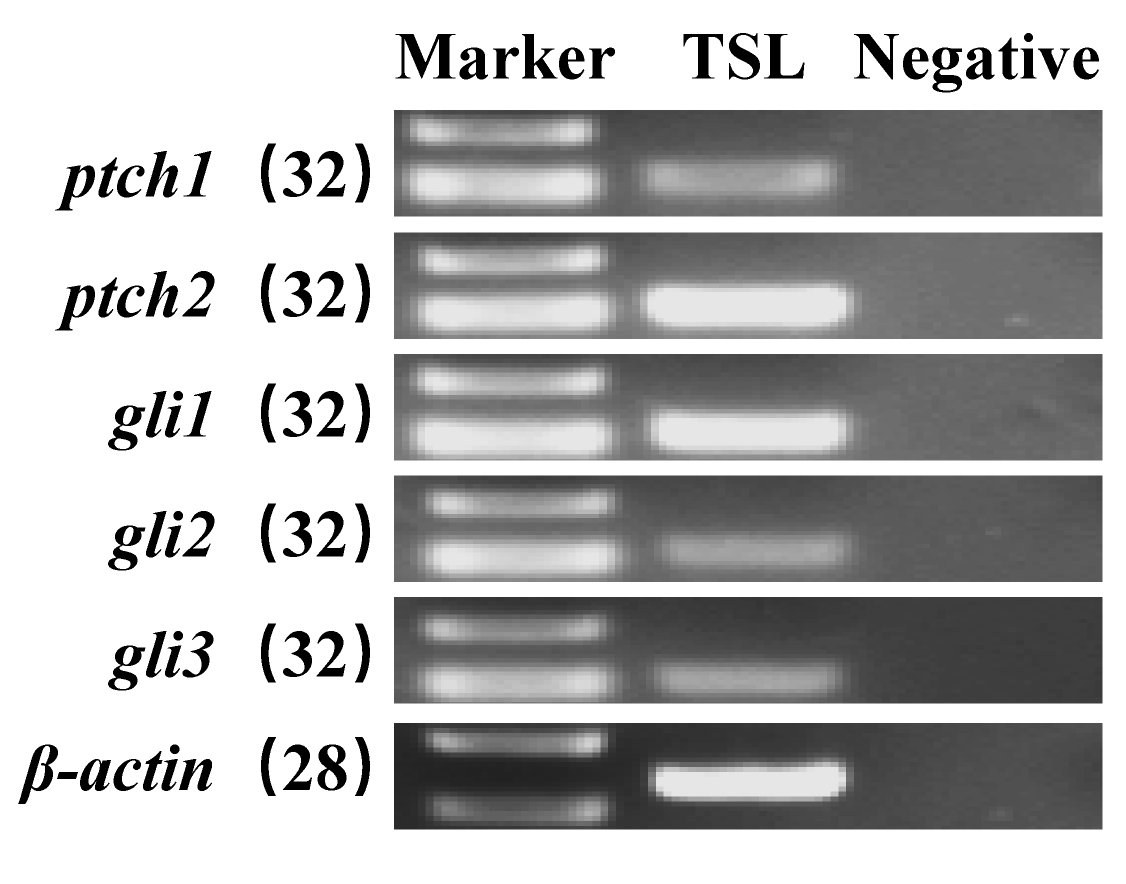

### Supplemental Figure S4

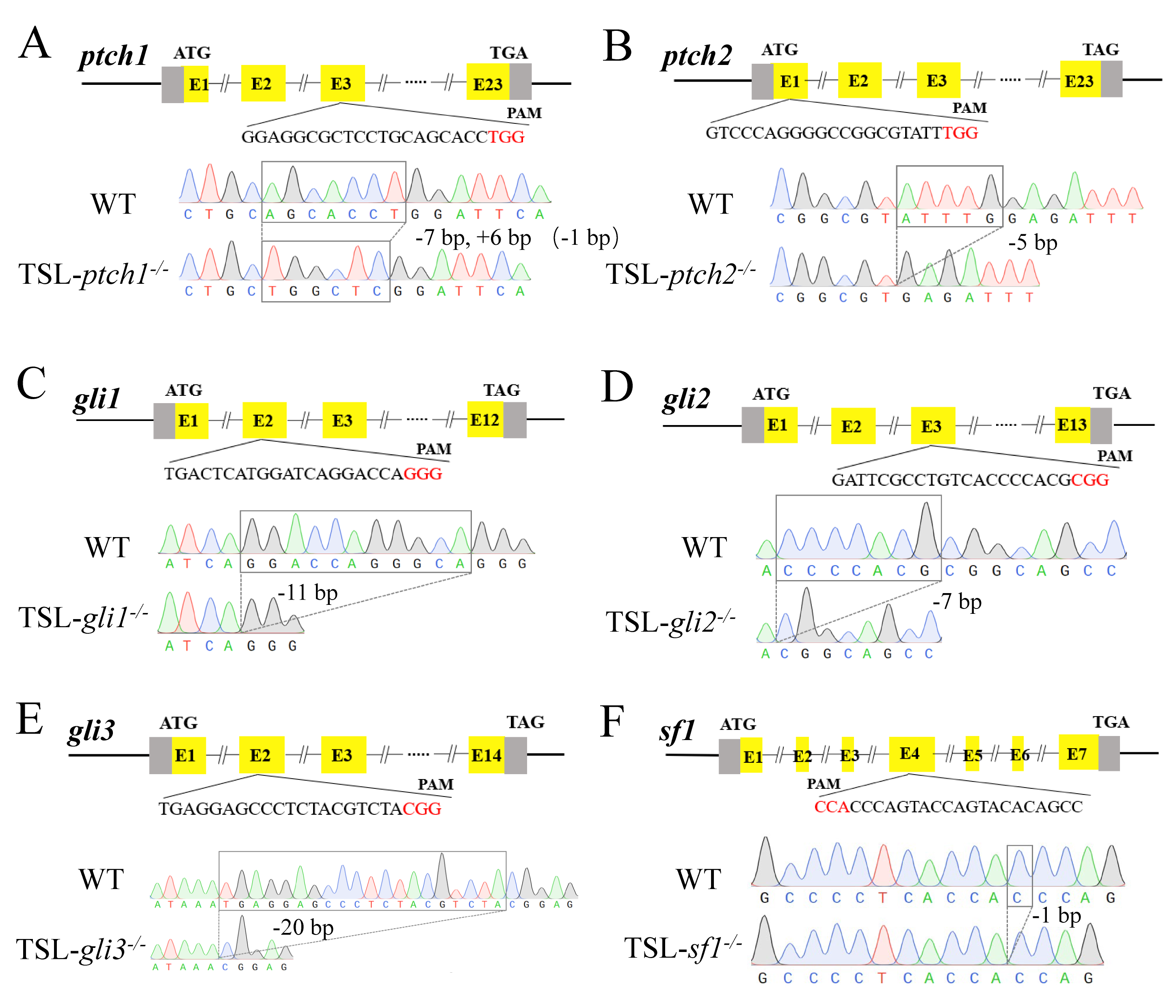

### Supplemental Figure S5

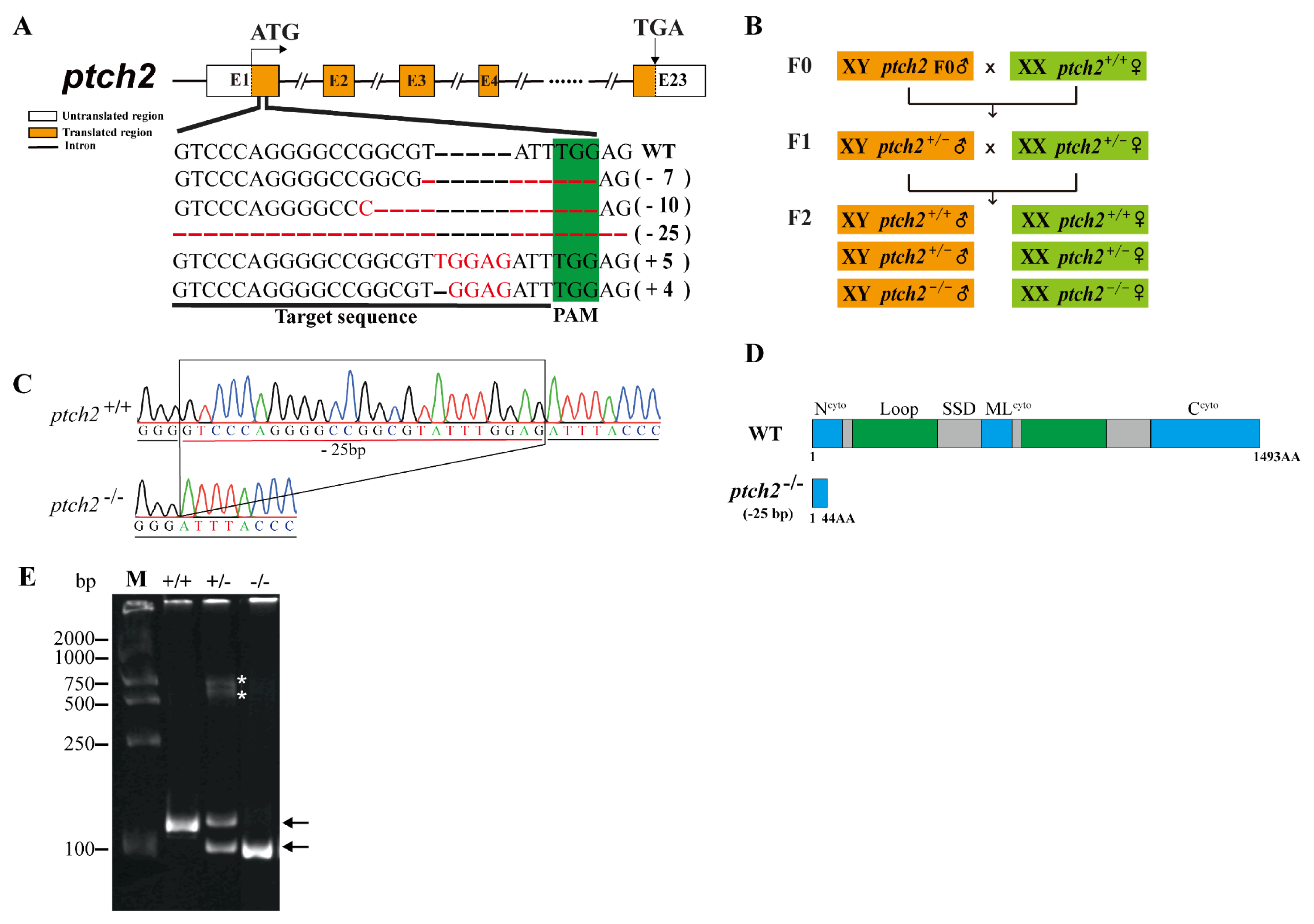

### Supplemental Figure S6

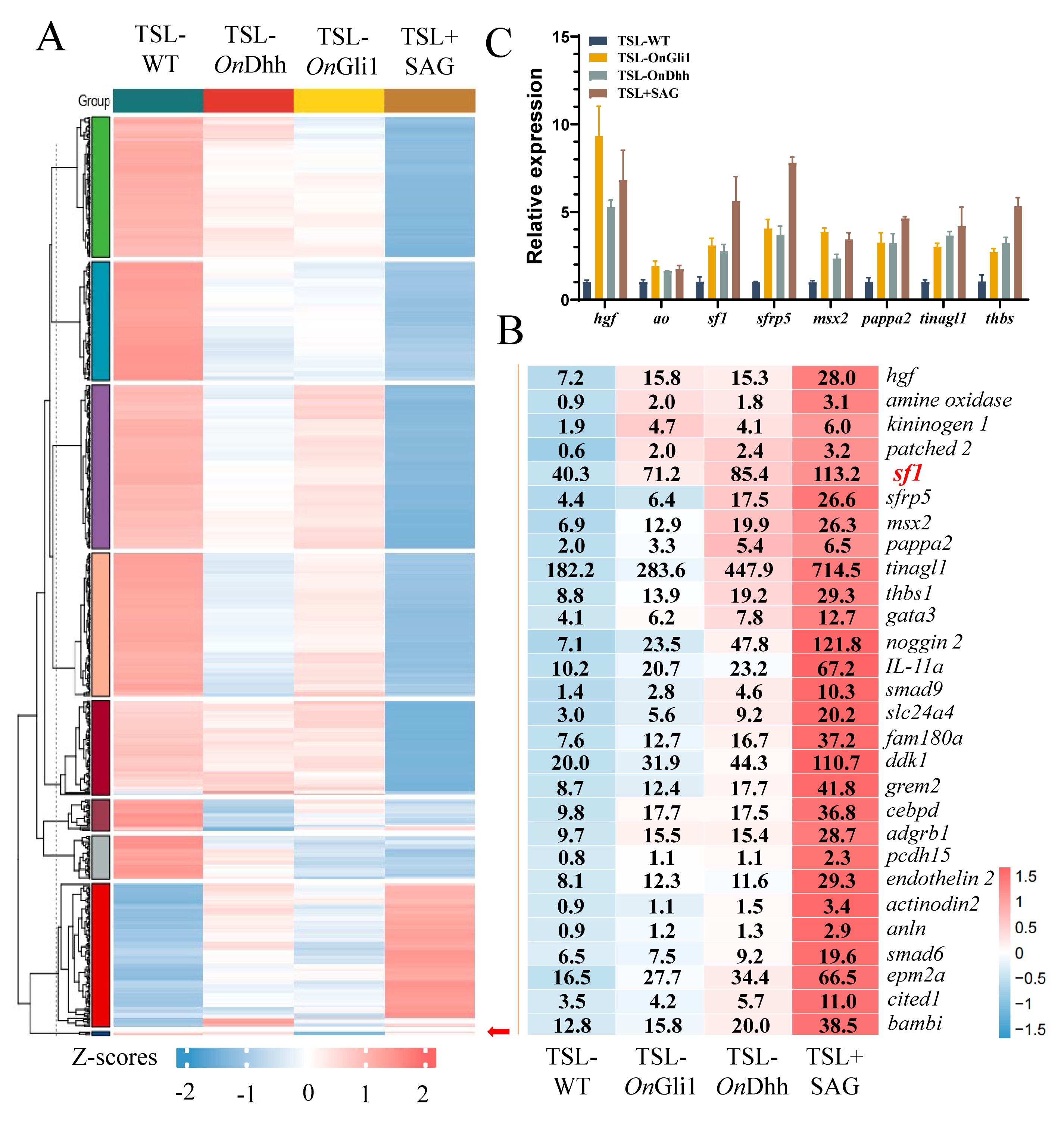

### Supplemental Figure S7

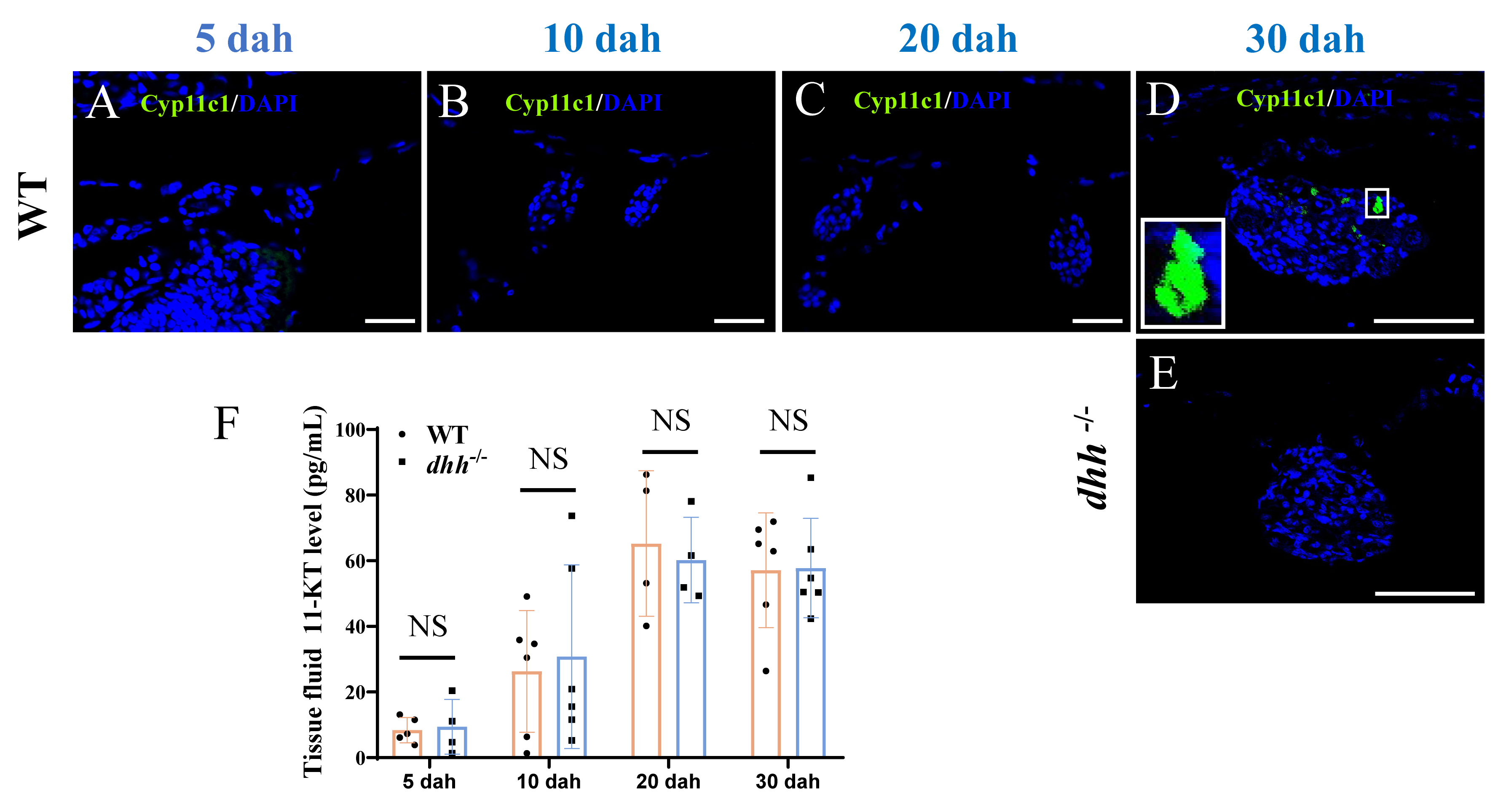

### Supplemental Figure S8

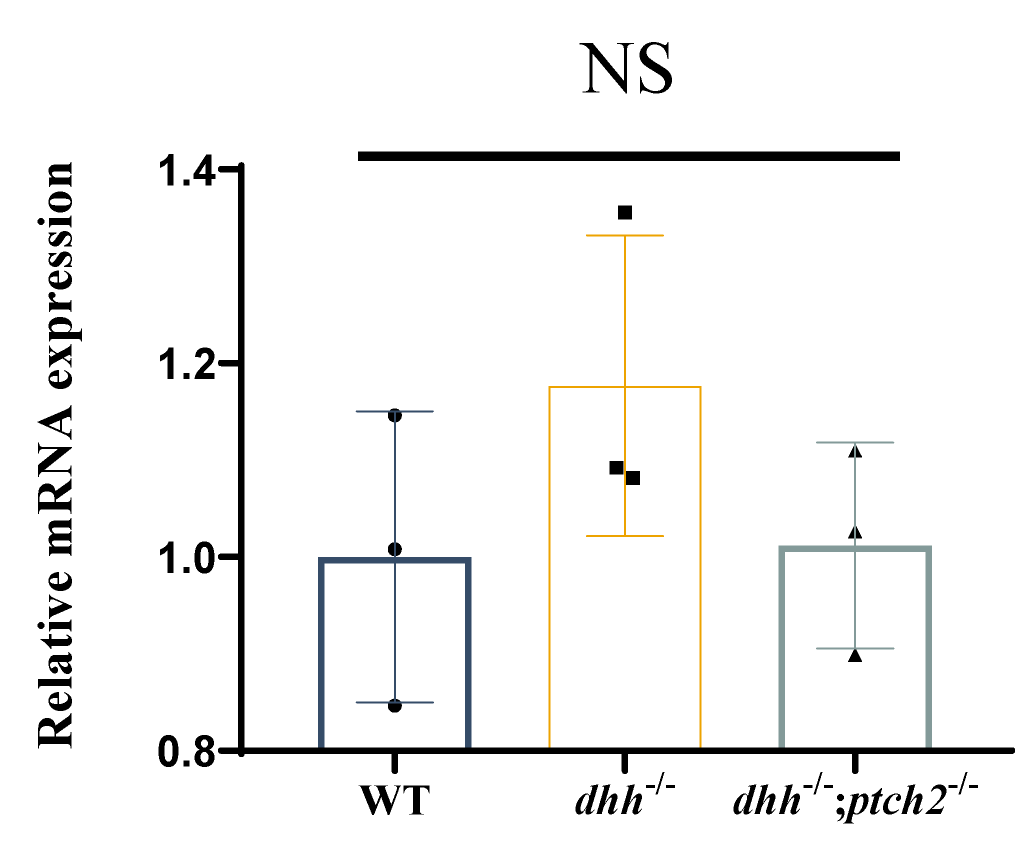
