## Supplemental Table S1 for "Desert Hedgehog mediates stem Leydig cell differentiation through Ptch2/Gli1/Sf1 signaling axis"

| Primer name | Sequence (5'-3') | Purpose |
|---|---|---|
| gRNA-*dhh* -F | TAATACGACTCACTATAGCGGGCCCGGTCCGCA TCCCGTTTTAGAGCTAGAAATAGC | CRISPR/Cas9 in Nile tilapia |
| gRNA-*ptch2* -F | TAATACGACTCACTATAGTCCCAGGGGCCGGCG TATTGTTTTAGAGCTAGAAATAGC | |
| gRNA-R | AGCACCGACTCGGTGCCAC | |
| TSL-*ptch1* -KO-R1 | GGTGCTGCAGGAGCGCCTCCCGACAGCTCCAAG GACCCGGGAG | CRISPR/Cas9 in TSL |
| TSL-*ptch1* -KO-F1 | GGAGGCGCTCCTGCAGCACCGTTTTAGAGCTAG AAATAGC | |
| TSL-*ptch2* -KO-R1 | AATACGCCGGCCCCTGGGACCGACAGCTCCAAG GACCCGGGAG | |
| TSL-*ptch2* -KO-F1 | GTCCCAGGGGCCGGCGTATTGTTTTAGAGCTAG AAATAGC | |
| TSL-*gli1* -KO-R1 | TGGTCCTGATCCATGAGTCACGACAGCTCCAAG GACCCGGGAG | |
| TSL-*gli1* -KO-F1 | TGACTCATGGATCAGGACCAGTTTTAGAGCTAG AAATAGC | |
| TSL-*gli2* -KO-R1 | CGTGGGGTGACAGGCGAATCCGACAGCTCCAAG GACCCGGGAG | |
| TSL-*gli2* -KO-F1 | GATTCGCCTGTCACCCCACGGTTTTAGAGCTAGA AATAGC | |
| TSL-*gli3* -KO-R1 | TAGACGTAGAGGGCTCCTCACGACAGCTCCAAG GACCCGGGAG | |
| TSL-*gli3* -KO-F1 | TGAGGAGCCCTCTACGTCTAGTTTTAGAGCTAG AAATAGC | |
| TSL-*sf1* -KO-R1 | CCCAGTACCAGTACACAGCCCGACAGCTCCAAG GACCCGGGAG | |
| TSL-*sf1* -KO-F1 | GGCTGTGTACTGGTACTGGGGTTTAGAGCTAG AAATAGC | |
| Mlu-F | CTTGACGAGTTCTTCTGAACGCGTCTCGAGCCTC TAGA | |
| sal-R | CGCCATATTGAATTGGCGGTCGACTGGCGTAAT AGCCAAC | |
| *dhh* -PAGE-F | TGGCACAGCTCACCCTGGTT | Mutant screening |
| *dhh* -PAGE-R | AAACTGCTTGAGGCGCATGG | |
| *dhh* -test-F | ACGTGCCCCGGAACTTTATT | |
| *dhh* -test-R | TAGTCATGAAGCGGTCAGCG | |
| *ptch1* -PAGE-F | CTGATGATCCAGACGCCGC | |
| *ptch1* -PAGE-R | CTCGAACAGAGGCAGTTATGG | |
| *ptch1* -test-F | CAATTCTGGCCCCTAACCCTT | |
| *ptch1* -test-R | GCTCTAACAGGGCTACACTCA | |
| *ptch2* -PAGE-F | GATTCTGTGTCGCTTCGCTCT | |
| *ptch2* -PAGE-R | AGTAGGTCTGGGTTCGCAGG | |
| *ptch2* -test-F | ATCGGGACGAATGAAGACGG | |
| *ptch2* -test-R | GACTCCGGTTTGATAGCTGC | |
| *gli1* -PAGE-F | CTGCCACGCTATGTATAATCCCA | |
| *gli1* -PAGE-R | GCCCATCATACCGTGGGGTG | |
| *gli1* -test-F | ATTCCTTGTCACAGTTCCCTTTG | |
| *gli1* -test-R | AGAGTAATCCACTCGCCTGCT | |
| *gli2* -PAGE-F | GGGCAGTCCAGTCATCTCAG | |
| *gli2* -PAGE-R | GACATACGGATGTGGAGGGC | |
| *gli2* -test-F | GGGGAGGGGAAAGGGTGTAA | |
| *gli2* -test-R | CTCCACCGCTCACTGACATT | |
| *gli3* -PAGE-F | AACGCCATCAGCTCTCAGG | |
| *gli3* -PAGE-R | GAACCGTGCTGCTCCTTCTT | |
| *gli3* -test-F | CTCATATCCAGGGGCTGAAGG | |
| *gli3* -test-R | GAGTCTAGGTAGCCGTTGCG | |

| | | |
|---|---|---|
| *sf1* -PAGE-F | CTCTACGGACCCCCGTCTTT | |
| *sf1* -PAGE-R | CTCGGCTTTAATTGCCCTTCC | |
| *sf1* -test-F | AGGGCCTTGAAACAGCAGAA | |
| *sf1* -test-R | ATGCGACACATGAGGCTGAA | |
| *ptch1* -ISH-F | AAAAAGCGGACATTGGGCAC | |
| *ptch1* -ISH-R | GCCAAAAAGGGAAGCACCTG | |
| *ptch2* -ISH-F | GTCATCCTGATCGCTTCCGT | |
| *ptch2* -ISH-R | AGTGGCTTCACCACCGTAAG | |
| *gli1* -ISH-F | GGCCCCTGCTTGGATCAATA | *In situ hybridization* |
| *gli1* -ISH-R | CAATCGCACCTCCAAGACCT | |
| *gli2* -ISH-F | GGCGGTACCATTATGAGCCA | |
| *gli2* -ISH-R | AAAGGTGCCCATAGGAGCTG | |
| *gli3* -ISH-F | GCAACGGCTACCTAGACTCG | |
| *gli3* -ISH-R | GTCGATCGCCTGCAGATACT | |
| 8xGLI-PGL4.23-F1 | CTAGGACCACCCAGACCACCCAGACCACCCAGA | |
| 8xGLI-PGL4.23-R1 | AGCTTGGGTGGTCTGGGTGGTCTGGGTGGTCTG | |
| NT-Flag-Dhh-pcDNA3.1-F1 | CTAGCTAGCATGAAGCAGTTCTGGTGGGC | |
| NT-Flag-Dhh-pcDNA3.1-R1 | CCGCTCGAGTTATTTATCATCATCATCTTTATAA | |
| NT-Flag-Gli1-pcDNA3.1-F1 | TTTAAACTTAAGCTTGGTACATGGATTATAAAG | |
| NT-Flag-Gli1-pcDNA3.1-R1 | TGCGCATGTGAACCACCAGC | |
| NT-Flag-Gli1-pcDNA3.1-F2 | GCTGGTGGTTCACATGCGCA | |
| NT-Flag-Gli1-pcDNA3.1-R2 | AACCCTCCTCCAGGACACGG | |
| NT-Flag-Gli1-pcDNA3.1-F3 | CCGTGTCCTGGAGGAGGGTT | Luciferase |
| NT-Flag-Gli1-pcDNA3.1-R3 | GTTTGCTGCAGAAGCATGCG | |
| NT-Flag-Gli1-pcDNA3.1-F4 | CGCATGCTTCTGCAGCAAAC | |
| NT-Flag-Gli1-pcDNA3.1-R4 | TCCACCACACTGGACTAGTGCTAAGACAGTGTA | |
| NT-*sf1* -pGL3-F1 | CGAGATCTGCGATCTAAGTATTACAAGTTAGAC TTTTGTA | |
| NT-*sf1* -pGL3-R1 | GATTGCACCTTCAGTGCTCCTGGC | |
| NT-*sf1* -pGL3-F2 | GCCAGGAGCACTGAAGGTGCAATC | |
| NT-*sf1* -pGL3-R2 | CCAACAGTACCGGAATGCCATTGAGCAGCAGAC AGCAGGC | |
| *sf1* -qPCR-F | TGTCCCTTCTGTCGGTTCCA | |
| *sf1* -qPCR-R | TCTCTTGTACATGGGGCCAA | |
| *tinagl1* -qPCR-F | CTGGGCACACAAAGACCATC | |
| *tinagl1* -qPCR-R | ATCGAGCGGTTCGTGAATCT | |
| *hgf* -qPCR-F | GTGCTTTACCACCGACCCTC | |
| *hgf* -qPCR-R | ACTTTCTGTGTGGTCCATTGGT | |
| *msx2* -qPCR-F | CAGCGAAGACAGTCCACAGTT | |
| *msx2* -qPCR-R | TAGTAAAGGGCGTGCGAGGT | Real-time PCR |
| *amine oxidase* -qPCR-F | TGTTATAGTGGTCGGTGCGG | |
| *amine oxidase* -qPCR-R | GCACAGTGAAGGTACGACCA | |
| *thbs1* -qPCR-F | CACAATGGCATCGTTCGCAA | |
| *thbs1* -qPCR-R | CTCCATCGGGAACAGTAGCA | |
| *pappa2* -qPCR-F | CACGTTCAGCCTGGAGCTTT | |
| *pappa2* -qPCR-R | TGCCTACTGACCAACCCTTCT | |
| *sfrp5* -qPCR-F | TCCGGTGATGGAGACCTACG | |
| *sfrp5* -qPCR-R | CACCTTTGACACTGGTGGCT | |

forward primer; R, reverse primer
